# Designing Novel Subunit Vaccines against Herpes Simplex Virus-1 using Reverse Vaccinology Approach

**DOI:** 10.1101/2020.01.10.901678

**Authors:** Bishajit Sarkar, Md. Asad Ullah

**Author notes:** Corresponding author: Bishajit Sarkar Email of the corresponding author.

## Abstract

Herpes Simplex Virus (HSV) is an infectious virus that is responsible for various types of orofacial and genital infections. Two types of HSV viruses, HSV-1 and HSV-2, are the most dangerous HSV viruses. Every year, millions of people get infected with this menacing virus, however, no satisfactory treatment or vaccine has not yet been discovered to fight against HSV. Although there are some anti-viral therapies, however, studies have showed that such anti-viral therapies may also fail to provide good impact. In this study, a possible subunit vaccine against HSV-1, strain-17, was designed using the tools and reverse vaccinology and bioinformatics. Three potential antigenic envelope glycoproteins were selected from nine envelope glycoproteins, for possible vaccine construction. Potential epitopes capable of inducing high immunogenic response and at the same time have non-allergenicity and conservancy across other strains and species, were selected by some robust analysis, for vaccine construction. Finally, three possible vaccines were designed. Each of the vaccine construct differ from each other only in their adjuvant sequences and based on molecular docking analysis, one best vaccine construct was selected for molecular dynamics simulation study and in silico codon adaptation. The experiment showed that the selected best vaccine should be good candidate against HPV-1, strain-17. However, wet lab study should be conducted on the suggested vaccine(s) for confirming their potentiality, safety and efficacy.

## 1. Introduction

Herpes Simplex Virus (HSV) is a member of the herpesviridae family. HSV is a double stranded DNA containing virus with an icosapentahedral capsid. The HSV-1 and HSV-2 are the two most dangerous types of herpes. HSV-1 mainly causes the orofacial infections and HSV-2 usually causes genital infections [1, 2]. The HSV-1 virus is transmitted from individual to individual primarily by oral-oral contact and cause a disease which is known as cold sores. HSV-1 infection is wide-spread throughout the world and in many cases, the infection may remain latent or undetected throughout the whole life of the infected person. Sometimes, the infection may lead to much dangerous diseases like the encephalitis. On the other hand, the HSV-2 is mainly a sexually transmitted virus and therefore, it mainly causes the genital herpes. According to a study, the total number of new HSV-1 cases in 2012 was estimated to be 118 million [3, 4]. Moreover, the global incidence of HSV-2 was estimated to be about 417 million in 2012 [5]. Sometimes, the incidence of HSV infection leads to other diseases like the Alzheimer’s disease and liver failure [6, 7]. Currently, the treatment of HSV is carried out using many antiviral medications like acyclovir, valacyclovir, famciclovir etc. However, in many cases, such antiviral medications have failed to provide complete resistant to HSV and reduce the mortality rate of the HSV infected patients. Moreover, studies have proved that the viral resistance of Herpes virus to such medications (like the acyclovir) may also occur and such incidence can make the treatments more complicated [8, 9, 10, 11]. Although researchers are working on to develop an effective vaccine against the Herpes virus like subunit vaccines, inactivated virion vaccines, genetically attenuated vaccines etc., to date, no satisfactory vaccine has been successfully entered the market [12]. In this study, the reverse vaccinology and bioinformatics approach was used to develop an effective Herpes vaccine against the HSV-1, strain-17 (**Figure 01**). Reverse vaccinology is defined as the process of vaccine development where the novel antigens of a virus or organism are identified by analyzing the genomic information of that particular organism or a virus. In reverse vaccinology, various tools of bioinformatics are used for identifying the novel antigens useful for vaccine development by dissecting the genome as well as for studying the genetic makeup of a pathogen. This approach of vaccine development also helps the scientists to understand which antigenic segments of a virus or pathogen should be given emphasis during the vaccine development. This method is a quick easy and cost-effective way to design vaccine. Reverse vaccinology is successfully used for developing vaccines for many viruses like the Zika virus, Chikungunya virus etc. [13, 14].

## 2. Materials and methods

The experiment focuses on developing of vaccines against the Human Herpes Simplex Virus-1 (strain-17).

### 2.1. Strain Identification and Selection

The Herpes Simplex Virus-1 (strain-17) was identified and selected by analyzing and reviewing different entries of the online server of National Center for Biotechnology Information or NCBI (https://www.ncbi.nlm.nih.gov/).

### 2.2. Retrieval of the Protein Sequences

Nine viral proteins: envelope glycoprotein M (accession number: P04288), envelope glycoprotein H (accession number: P06477), envelope glycoprotein E (accession number: P04488), envelope glycoprotein I (accession number: P06487), envelope glycoprotein B (accession number: P10211), envelope glycoprotein L (accession number: P10185), envelope glycoprotein D (accession number: Q69091), envelope glycoprotein K (accession number: P68331) and envelope glycoprotein C (accession number: P10228) were retrieved from the UniProt Knowledgebase (UniProtKB) tool of the online server UniProt (https://www.uniprot.org/).

### 2.3. Antigenicity Prediction and Physicochemical Property Analysis of the Protein Sequences

The antigenicity of the protein sequences were predicted by the online server, VaxiJen v2.0 (http://www.ddg-pharmfac.net/vaxijen/VaxiJen/VaxiJen.htm), keeping the threshold at 0.4 and tumor model was used [15, 16, 17]. Only the antigenic proteins were selected for further analysis. The various physicochemical properties of the selected antigenic protein sequences were determined by ExPASy’s online tool ProtParam (https://web.expasy.org/protparam/) [18].

### 2.4. T-cell and B-cell Epitope Prediction

The T-cell and B-cell epitopes of the selected protein sequences were predicted using online epitope prediction server Immune Epitope Database or IEDB (https://www.iedb.org/). The IEDB database contains huge amount of experimental data on T-cell epitopes as well as antibodies. These data are collected from various studies that are conducted on human, non-human primates and other animals. It is a server that allows robust analysis on various epitopes by exploiting various tools: population coverage, conservation across antigens and clusters with similar sequences [19]. The MHC class-I restricted CD8+ cytotoxic T-lymphocyte (CTL) epitopes of the selected sequences were obtained using NetMHCpan EL 4.0 prediction method for HLA-A*11-01 allele. The MHC class-II restricted CD4+ helper T-lymphocyte (HTL) epitopes were obtained for HLA DRB1*04-01 allele using Sturniolo prediction method. The top ten MHC class-I were selected based on their percentile scores and antigenicity scores (AS). Moreover, MHC class-II epitope prediction generated several epitopes with similar AS and percentile scores. However, 2 epitopes were taken from each of the top five categories. On the other hand, five random B-cell lymphocyte epitopes (BCL) were selected based on their length (the epitope sequences that had ten amino acids or above were selected) and obtained using Bipipered linear epitope prediction method, keeping the parameters default.

### 2.5. Transmembrane Topology and Antigenicity Prediction of the Selected Epitopes

The transmembrane topology of the selected epitopes were determined using the transmembrane topology of protein helices determinant, TMHMM v2.0 server (http://www.cbs.dtu.dk/services/TMHMM/). The server predicts whether the epitope would be transmembrane or it would remain inside or outside of the membrane [20]. The antigenicity of the selected epitopes were predicted using VaxiJen v2.0 (http://www.ddg-pharmfac.net/vaxijen/VaxiJen/VaxiJen.htm), using the tumor model and threshold of 0.4.

### 2.6. Allergenicity and Toxicity Prediction of the Epitopes

The allergenicity of the selected epitopes were predicted using two online tools, AllerTOP v2.0 (https://www.ddg-pharmfac.net/AllerTOP/) and AllergenFP v1.0 (http://ddg-pharmfac.net/AllergenFP/). However, the results predicted by AllerTOP were given priority since the server has better accuracy of 88.7% than AllergenFP server (87.9%) [21, 22]. The toxicity prediction of the selected epitopes were carried out using ToxinPred server (http://crdd.osdd.net/raghava/toxinpred/), using SVM (Swiss-Prot) based method, keeping all the parameters default.

### 2.7. Conservancy Prediction of the Selected Epitopes

The conservancy analysis of the selected epitopes were performed using the ‘epitope conservancy analysis’ tool of IEDB server (https://www.iedb.org/conservancy/) [19]. The sequence identity threshold was kept ‘>=50’. For the conservancy analysis of the selected epitopes of the Herpes Simplex Virus-1 strain-17, the envelope glycoprotein E, envelope glycoprotein B and envelope glycoprotein D of Human Herpes Simplex Virus-1 strain-F and Human Herpes Simplex Virus-2 strain-HG52 were used for comparison along with the proteins of the Human Herpes Simplex Virus-1 strain-17 itself (UniProt accession numbers: P89475, Q703F0, P04488, P08666, P06436, P10211, Q69467, Q05059 and Q69091). This ensures that, the epitope sequences are conserved across the species as well as strains. Based on the antigenicity, allergenicity, toxicity and conservancy analysis, the best ligands were selected for the further analysis and vaccine construction. The epitopes that showed antigenicity, non-allergenicity, non-toxicity and high (more than 90%) conservancy and more than 50% minimum identity, were considered as the best epitopes. For B-cell epitope selection, only the antigenic and non-allergenic epitopes were taken for further analysis.

### 2.8. Cluster Analysis of the MHC Alleles

Cluster analysis of the MHC alleles helps to identify the alleles of the MHC class-I and class-II molecules that have similar binding specificities. The cluster analysis of the MHC alleles were carried out by online tool MHCcluster 2.0 (http://www.cbs.dtu.dk/services/MHCcluster/) [23]. During the analysis, the number of peptides to be included was kept 50,000, the number of bootstrap calculations were kept 100 and all the HLA supertype representatives (MHC class-I) and HLA-DR representatives (MHC class-II) were selected. For analyzing the MHC class-I alleles, the NetMHCpan-2.8 prediction method was used. The output of the server generated results in the form of MHC specificity tree and MHC specificity heat-map.

### 2.9. Generation of the 3D Structures of the Selected Epitopes

The 3D structures of the selected best epitopes were generated using online 3D generating tool PEP-FOLD3 (http://bioserv.rpbs.univ-paris-diderot.fr/services/PEP-FOLD3/). PEP-FOLD3 is a online tool for generating de novo peptide 3D structure [24, 25, 26].

### 2.10. Molecular Docking of the Selected Epitopes

The molecular docking of the selected epitopes with the MHC class-I and class-II proteins were carried out by online docking tool PatchDock (https://bioinfo3d.cs.tau.ac.il/PatchDock/php.php). PatchDock tool divides the Connolly dot surface representation of the molecules into concave, convex and flat patches using its various algorithms. After that the complementary patches are matched for generating potential candidate transformations. Next, each of the candidate transformations is evaluated by a scoring function and later, an RMSD (root mean square deviation) clustering is applied to the candidate solutions for discarding the redundant solutions. The top score solutions are made the top ranked solutions by the server. After docking by PatchDock, the docking results were refined and re-scored by FireDock server (http://bioinfo3d.cs.tau.ac.il/FireDock/php.php). The FireDock server generates global energies upon the refinement of the best solutions from the PatchDock server and ranks them based on the generated global energies and the lowest global energy is always appreciable and preferred [27, 28, 29, 30]. The molecular docking experiments were carried out using the HLA-A*11-01 allele (PDB ID: 5WJL) and HLA DRB1*04-01 (PDB ID: 5JLZ) as receptors and the ligands were the best selected MHC-I and MHC-II epitopes, respectively. The receptors were downloaded from the RCSB-Protein Data Bank (https://www.rcsb.org/) server. The best results were visualized using Discovery Studio Visualizer [31].

### 2.11. Vaccine Construction

Three possible vaccines were constructed against the Herpes Simplex Virus-1, strain-17. The predicted CTL, HTL and BCL epitopes were conjugated together for constructing the vaccines. All the vaccines were generated maintaining the sequence: adjuvant, PADRE sequence, CTL epitopes, HTL epitopes and BCL epitopes. Three different adjuvant sequences were used for constructing three different vaccines: beta defensin, L7/L12 ribosomal protein and HABA protein (*M. tuberculosis*, accession number: AGV15514.1). Beta-defensin adjuvants induce the activation of the toll like receptors (TLRs): 1, 2 and 4, where beta-defensin acts as agonist. The L7/L12 ribosomal protein and HABA protein activate the TLR-4. During the vaccine construction, various linkers were used: EAAAK linkers were used to conjugate the adjuvant and PADRE sequence, GGGS linkers were used to attach the PADRE sequence with the CTL epitopes and the CTL epitopes with the other CTL epitopes, GPGPG linkers were used to connect the CTL with HTL epitopes and also the HTL epitopes among themselves. The KK linkers were used for conjugating the HTL and BCL epitopes as well as the BLC epitopes among themselves [32, 33, 34, 35, 36, 37, 38]. Studies have proved that the PADRE sequence improves the CTL response of the vaccines that contain it [39]. Total three vaccines were constructed in the experiment.

### 2.12. Antigenicity, Allergenicity and Physicochemical Property Analysis

The antigenicity of the constructed vaccines were determined by the online server VaxiJen v2.0 (http://www.ddg-pharmfac.net/vaxijen/VaxiJen/VaxiJen.htm). The threshold of the prediction was kept at 0.4. AlgPred (http://crdd.osdd.net/raghava/algpred/) and AllerTop v2.0 (https://www.ddg-pharmfac.net/AllerTOP/) were used for the prediction of the allergenicity of the vaccine constructs. The AlgPred server predicts the possible allergens based on similarity of known epitope of any of the known region of the protein [40]. MEME/MAST motif prediction approach is used in the allergenicity prediction of the vaccines by AlgPred. Moreover, various physicochemical properties of the vaccines were examined by the online server ProtParam (https://web.expasy.org/protparam/).

### 2.13. Secondary and Tertiary Structure Prediction of the Vaccine Constructs

The secondary structures of the vaccine constructs were generated using online tool PRISPRED (http://bioinf.cs.ucl.ac.uk/psipred/). PRISPRED is a simple secondary structure generator which can be used to predict the transmembrane topology, transmembrane helix, fold and domain recognition etc. along with the secondary structure prediction [41, 42]. The PRISPRED 4.0 prediction method was used to predict the secondary structures of the vaccine constructs. The β-sheet structure of the vaccines were determined by another online tool, NetTurnP v1.0 (http://www.cbs.dtu.dk/services/NetTurnP/) [43]. The tertiary or 3D structures of the vaccines were generated using online tool RaptorX (http://raptorx.uchicago.edu/) server. The server is a fully annotated tool for the prediction of protein structure, the property and contact prediction, sequence alignment etc. [44, 45, 46].

## 2.14. 3D Structure Refinement and Validation

The 3D structures of the constructed vaccines were refined using online refinement tool, 3Drefine (http://sysbio.rnet.missouri.edu/3Drefine/). The server is a quick, easy and efficient tool for protein structure refinement [47]. For each of the vaccine, the refined model 1 was downloaded for validation. The refined vaccine proteins were then validated by analyzing the Ramachandran plots which were generated using the online tool, PROCHECK (https://servicesn.mbi.ucla.edu/PROCHECK/) [48, 49].

### 2.15. Vaccine Protein Disulfide Engineering

The vaccine protein disulfide engineering was carried out by online tool Disulfide by Design 2 v12.2 (http://cptweb.cpt.wayne.edu/DbD2/). The server predicts the possible sites within a protein structure which have the greater possibility of undergoing disulfide bond formation [50]. When engineering the disulfide bonds, the intra-chain, inter-chain and C_β_ for glycine residue were selected. The χ^3^ Angle was kept −87° or +97° ± 5 and C_α_-C_β_-S_γ_ Angle was kept 114.6° ±10.

### 2.16. Protein-Protein Docking

In protein-protein docking, the constructed Herpes Simplex Virus-1, strain-17 virus vaccines were analyzed by docking against various MHC alleles and toll like receptor (TLR). One best vaccine was selected based on their performances in the docking experiment. When viral infections occur, the viral particles are recognized by the MHC complex as antigens. The various segments of the MHC molecules are encoded by different alleles. For this reason, the vaccines should have good binding affinity with these MHC portions that are encoded by different alleles [51]. All the vaccine constructs were docked against the selected MHC alleles to test their binding affinity. In this experiment, the vaccines constructs were docked against DRB1*0101 (PDB ID: 2FSE), DRB3*0202 (PDB ID: 1A6A), DRB5*0101 (PDB ID: 1H15), DRB3*0101 (PDB ID: 2Q6W),

DRB1*0401 (PDB ID: 2SEB), and DRB1*0301 (PDB ID: 3C5J) alleles. Moreover, studies have proved that TLR-8 that are present on the immune cells, are responsible for mediating the immune responses against the RNA viruses and TLR-3 of the immune cells mediates immune responses against the DNA viruses [52, 53]. The Herpes virus is a DNA virus 9540. For this reason, the vaccine constructs of Herpes virus were also docked against TLR-3 (PDB ID: 2A0Z). The protein-protein docking was carried out using various online docking tools. The docking was carried out three times by three different online servers to improve the accuracy of the docking. First, the docking was carried out by ClusPro 2.0 (https://cluspro.bu.edu/login.php). The server ranks the clusters of docked complexes based on their center and lowest energy scores. However, these scores do not represent the actual binding affinity of the proteins with their targets [55, 56, 57]. The bonding affinity (ΔG in kcal mol^-1^) of docked complexes were generated by PRODIGY tool of HADDOCK webserver (https://haddock.science.uu.nl/). The lower the binding energy generated by the server, the higher the binding binding affinity [58, 59, 60]. Moreover, the docking was again performed by PatchDock (https://bioinfo3d.cs.tau.ac.il/PatchDock/php.php) and later refined and re-scored by FireDock server (http://bioinfo3d.cs.tau.ac.il/FireDock/php.php). The FireDock server ranks the docked complexes based on their global energy and the lower the score, the better the result. Later, the docking was performed using HawkDock server (http://cadd.zju.edu.cn/hawkdock/). The Molecular Mechanics/Generalized Born Surface Area (MM-GBSA) study was also carried out using HawkDock server. According to the server, the lower score and lower energy corresponds to better scores [61, 62, 63, 64]. The HawkDock server generates several models of docked complex and ranks them by assigning HawkDock scores in the ascending order. For each of the vaccines and their respective targets, the score of model 1 was taken for analysis. Furthermore, the model 1 of every complex was analyzed for MM-GBSA study. From the docking experiment, one best vaccine was selected for further analysis. The docked structures were visualized by PyMol tool [65].

### 2.17. Molecular Dynamic Simulation

The molecular dynamics simulation study was performed on the best selected vaccine, HV-1. The study was carried out by the online server iMODS (http://imods.chaconlab.org/). iMODS can be used efficiently to investigate the structural dynamics of the protein complexes, since it is a quick, easy and user-friendly server. The server provides the values of deformability, B-factor (mobility profiles), eigenvalues, variance, co-variance map and elastic network, for a protein or docked protein-protein complex. The deformability depends on the ability to deform at each of its amino acid. The eigenvalue is related with the energy that is required to deform the given structure and the lower eigenvalue corresponds to the easier the deformability of the complex. The eigenvalue also represents the motion stiffness of the protein complex. The server is a fast and easy tool for determining and measuring the protein flexibility [66, 67, 68, 69, 70]. For analysing the molecular dynamics simulation of the HV-1-TLR-3 docked complex was used. The docked PDB files were uploaded to the iMODS server and the results were displayed keeping all the parameters as default.

### 2.18. Codon Adaptation and In Silico Cloning

The codon adaptation and in silico cloning were carried out for the best selected vaccine protein, HV-1. For conducting these experiments, the vaccine protein was reverse transcribed to the possible DNA sequences. The DNA sequence should encode the target vaccine protein. Later, the reverse transcribed DNA sequences were adapted according to the desired organism, so that the cellular mechanisms of that specific organism could use the codons of the adapted DNA sequences efficiently and provide better production of the desired product. Codon adaptation is a necessary step of in silico cloning since the same amino acid can be encoded by different codons in different organisms, a phenomenon which is known as codon biasness. Moreover, the cellular mechanisms of an organism may be different from another organism and a codon for a specific amino acid may not work in another organism. For this reason, codon adaptation step is performed that can predict the suitable codon that can encode a specific amino acid in a specific organism [71, 72]. The codon adaptation of the selected vaccine protein was carried out by the Java Codon Adaptation Tool or JCat server (http://www.jcat.de/) [71]. Eukaryotic *E. coli* strain K12 was selected and rho-independent transcription terminators, prokaryotic ribosome binding sites and SgrA1 and SphI cleavage sites of restriction enzymes, were avoided. In the JCat server, the protein sequences were reverse translated to the optimized possible DNA sequences. The optimized DNA sequences were taken and SgrA1 and SphI restriction sites were conjugated at the N-terminal and C-terminal sites, respectively. Next, the SnapGene [73] restriction cloning module was used to insert the new adapted DNA sequences between SgrA1 and SphI of pET-19b vector.

## 3. Results

### 3.1. Identification, Selection and Retrieval of Viral Protein Sequences

The Herpes Simplex Virus, strain-17 was identified and selected from the NCBI (https://www.ncbi.nlm.nih.gov/). Nine envelope proteins from the viral structure was selected for the possible vaccine construction. These proteins were: envelope glycoprotein M (accession number: P04288), envelope glycoprotein H (accession number: P06477), envelope glycoprotein E (accession number: P04488), envelope glycoprotein I (accession number: P06487), envelope glycoprotein B (accession number: P10211), envelope glycoprotein L (accession number: P10185), envelope glycoprotein D (accession number: Q69091), envelope glycoprotein K (accession number: P68331) and envelope glycoprotein C (accession number: P10228). The protein sequences were retrieved from the online server, UniProt (https://www.uniprot.org/). The protein sequences in fasta format:

>sp|P04288|GM_HHV11 Envelope glycoprotein M OS=Human herpesvirus 1 (strain 17) OX=10299 GN=gM PE=1 SV=1 MGRPAPRGSPDSAPPTKGMTGARTAWWVWCVQVATFVVSAVCVTGLLVLASVFRARFPCFYATASSYAGVNSTAEVRGGVAVPLRLDTQSLVGTYVITAVLLLAVAVYAVVGAVTSRYDRALDAGRRLAAARMAMPHATLIAGNVCSWLLQITVLLLAHRISQLAHLVYVLHFACLVYFAAHFCTRGVLSGTYLRQVHGLMELAPTHHRVVGPARAVLTNALLLGVFLCTADAAVSLNTIAAFNFNFSAPGMLICLTVLFAILVVSLLLVVEGVLCHYVRVLVGPHLGAVAATGIVGLACEHYYTNGYYVVETQWPGAQTGVRVALALVAAFALGMAVLRCTRAYLYHRRHHTKFFMRMRDTRHRAHSALKRVRSSMRGSRDGRHRPAPGSPPGIPEYAEDPYAISYGGQLDRYGDSDGEPIYDEVADDQTDVLYAKIQHPRHLPDDDPIYDTVGGYDPEPAED PVYSTVRRW

>sp|P06477|GH_HHV11 Envelope glycoprotein H OS=Human herpesvirus 1 (strain 17) OX=10299 GN=gH PE=1 SV=1 MGNGLWFVGVIILGVAWGQVHDWTEQTDPWFLDGLGMDRMYWRDTNTGRLWLPNTPDPQKPPRGFLAPPDELNLTTASLPLLRWYEERFCFVLVTTAEFPRDPGQLLYIPKTYLLGRPPNASLPAPTTVEPTAQPPPSVAPLKGLLHNPAASVLLRSRAWVTFSAVPDPEALTFPRGDNVATASHPSGPRDTPPPRPPVGARRHPTTELDITHLHNASTTWLATRGLLRSPGRYVYFSPSASTWPVGIWTTGELVLGCDAALVRARYGREFMGLVISMHDSPPVEVMVVPAGQTLDRVGDPADENPPGALPGPPGGPRYRVFVLGSLTRADNGSALDALRRVGGYPEEGTNYAQFLSRAYAEFFSGDAGAEQGPRPPLFWRLTGLLATSGFAFVNAAHANGAVCLSDLLGFLAHSRALAGLAARGAAGCAADSVFFNVSVLDPTARLQLEARLQHLVAEILEREQSLALHALGYQLAFVLDSPSAYDAVAPSAAHLIDALYAEFLGGRVLTTPVVHRALFYASAVLRQPFLAGVPSAVQRERARRSLLIASALCTSDVAAATNADLRTALARADHQKTLFWLPDHFSPCAASLRFDLDESVFILDALAQATRSETPVEVLAQQTHGLASTLTRWAHYNALIRAFVPEASHRCGGQSANVEPRILVPITHNASYVVTHSPLPRGIGYKLTGVDVRRPLFLTYLTATCEGSTRDIESKRLVRTQNQRDLGLVGAVFMRYTPAGEVMSVLLVDTDNTQQQIAAGPTEGAPSVFSSDVPSTALLLFPNGTVIHLLAFDTQPVAAIAPGFLAASALGVVMITAALAGILKVLRTSVPFFWRRE

>sp|P04488|GE_HHV11 Envelope glycoprotein E OS=Human herpesvirus 1 (strain 17) OX=10299 GN=gE PE=1 SV=1 MDRGAVVGFLLGVCVVSCLAGTPKTSWRRVSVGEDVSLLPAPGPTGRGPTQKLLWAVEPLDGCGPLHPSWVSLMPPKQVPETVVDAACMRAPVPLAMAYAPPAPSATGGLRTDFV WQERAAVVNRSLVIHGVRETDSGLYTLSVGDIKDPARQVASVVLVVQPAPVPTPPPTPADYDEDDNDEGEDESLAGTPASGTPRLPPPPAPPRSWPSAPEVSHVRGVTVRMETPEAILFSPGETFSTNVSIHAIAHDDQTYSMDVVWLRFDVPTSCAEMRIYESCLYHPQLPECLSPADAPCAASTWTSRLAVRSYAGCSRTNPPPRCSAEAHMEPVPGLAWQAASVNLEFRDASPQHSGLYLCVVYVNDHIHAWGHITISTAAQYRNAVVEQPLPQRGADLAEPTHPHVGAPPHAPPTHGALRLGAVMGAALLLSALGLSVWACMTCWRRRAWRAVKSRASGKGPTYIRVADSELYADWSSDSEGERDQVPWLAPPERPDSPSTNGSGFEILSPTAPSVYPRSDGHQSRRQLTTFGSGRPDRRYSQASDSSVFW

>sp|P06487|GI_HHV11 Envelope glycoprotein I OS=Human herpesvirus 1 (strain 17) OX=10299 GN=gI PE=1 SV=1 MPCRPLQGLVLVGLWVCATSLVVRGPTVSLVSNSFVDAGALGPDGVVEEDLLILGELRFVGDQVPHTTYYDGGVELWHYPMGHKCPRVVHVVTVTACPRRPAVAFALCRATDSTHSPAYPTLELNLAQQPLLRVQRATRDYAGVYVLRVWVGDAPNASLFVLGMAIAAEGTLAYNGSAYGSCDPKLLPSSAPRLAPASVYQPAPNQASTPSTTTSTPSTTIPAPSTTIPAPQASTTPFPTGDPKPQPPGVNHEPPSNATRATRDSRYALTVTQIIQIAIPASIIALVFLGSCICFIHRCQRRYRRSRRPIYSPQMPTGISCAVNEAAMARLGAELKSHPSTPPKSRRRSSRTPMPSLTAIAEESEPAGAAGLPTPPVDPTTPTPTPPLLV

>sp|P10211|GB_HHV11 Envelope glycoprotein B OS=Human herpesvirus 1 (strain 17) OX=10299 GN=gB PE=1 SV=1 MRQGAPARGRRWFVVWALLGLTLGVLVASAAPSSPGTPGVAAATQAANGGPATPAPPAPGAPPTGDPKPKKNRKPKPPKPPRPAGDNATVAAGHATLREHLRDIKAENTDANFYV

CPPPTGATVVQFEQPRRCPTRPEGQNYTEGIAVVFKENIAPYKFKATMYYKDVTVSQVWFGHRYSQFMGIFEDRAPVPFEEVIDKINAKGVCRSTAKYVRNNLETTAFHRDDHETDMELKPANAATRTSRGWHTTDLKYNPSRVEAFHRYGTTVNCIVEEVDARSVYPYDEFVLATGDFVYMSPFYGYREGSHTEHTSYAADRFKQVDGFYARDLTTKARATAPTTRNLLTTPKFTVAWDWVPKRPSVCTMTKWQEVDEMLRSEYGGSFRFSSDAISTTFTTNLTEYPLSRVDLGDCIGKDARDAMDRIFARRYNATHIKVGQPQYYLANGGFLIAYQPLLSNTLAELYVREHLREQSRKPPNPTPPPPGASANASVERIKTTSSIEFARLQFTYNHIQRHVNDMLGRVAIAWCELQNHELTLWNEARKLNPNAIASATVGRRVSARMLGDVMAVSTCVPVAADNVIVQNSMRISSRPGACYSRPLVSFRYEDQGPLVEGQLGENNELRLTRDAIEPCTVGHRRYFTFGGGYVYFEEYAYSHQLSRADITTVSTFIDLNITMLEDHEFVPLEVYTRHEIKDSGLLDYTEVQRRNQLHDLRFADIDTVIHADANAAMFAGLGAFFEGMGDLGRAVGKVVMGIVGGVVSAVSGVSSFMSNPFGALAVGLLVLAGLAAAFFAFRYVMRLQSNPMKALYPLTTKELKNPTNPDASGEGEEGGDFDEAKLAEAREMIRYMALVSAMERTEHKAKKKGTSALLSAKVTDMVMRKRRNTNYTQVPNKDGDADEDDL

>sp|P10185|GL_HHV11 Envelope glycoprotein L OS=Human herpesvirus 1 (strain 17) OX=10299 GN=gL PE=1 SV=1 MGILGWVGLIAVGVLCVRGGLPSTEYVIRSRVAREVGDILKVPCVPLPSDDLDWRYETPSAINYALIDGIFLRYHCPGLDTVLWDRHAQKAYWVNPFLFVAGFLEDLSYPAFPANTQETETRLALYKEIRQALDSRKQAASHTPVKAGCVNFDYSRTRRCVGRQDLGPTNGTSGRTPVLPPDDEAGLQPKPLTTPPPIIATSDPTPRRDAATKSRRRRPHSRRL

>sp|Q69091|GD_HHV11 Envelope glycoprotein D OS=Human herpesvirus 1 (strain 17) OX=10299 GN=gD PE=1 SV=1 MGGAAARLGAVILFVVIVGLHGVRSKYALVDASLKMADPNRFRGKDLPVLDQLTDPPGVRRVYHIQAGLPDPFQPPSLPITVYYAVLERACRSVLLNAPSEAPQIVRGASEDVRKQPYNLTIAWFRMGGNCAIPITVMEYTECSYNKSLGACPIRTQPRWNYYDSFSAVSEDNLGFLMHAPAFETAGTYLRLVKINDWTEITQFILEHRAKGSCKYALPLRIPPSACLSPQAYQQGVTVDSIGMLPRFIPENQRTVAVYSLKIAGWHGPKAPYTSTLLPPELSETPNATQPELAPEDPEDSALLEDPVGTVAPQIPPNWHIPSIQDAATPYHPPATPNNMGLIAGAVGGSLLAALVICGIVYWMRRHTQKAPKRIRLPHIREDDQPSSHQPLFY

>sp|P68331|GK_HHV11 Envelope glycoprotein K OS=Human herpesvirus 1 (strain 17) OX=10299 GN=gK PE=1 SV=1 MLAVRSLQHLSTVVLITAYGLVLVWYTVFGASPLHRCIYAVRPTGTNNDTALVWMKMNQTLLFLGAPTHPPNGGWRNHAHICYANLIAGRVVPFQVPPDAMNRRIMNVHEAVNCLETLWYTRVRLVVVGWFLYLAFVALHQRRCMFGVVSPAHKMVAPATYLLNYAGRIVSSVFLQYPYTKITRLLCELSVQRQNLVQLFETDPVTFLYHRPAIGVIVGCELMLRFVAVGLIVGTAFISRGACAITYPLFLTITTWCFVSTIGLTELYCILRRGPAPKNADKAAAPGRSKGLSGVCGRCCSIILSGIAVRLCYIAVVAGVVLVALHYEQEIQRRLFDV

>sp|P10228|GC_HHV11 Envelope glycoprotein C OS=Human herpesvirus 1 (strain 17) OX=10299 GN=gC PE=1 SV=1 MAPGRVGLAVVLWSLLWLGAGVSGGSETASTGPTITAGAVTNASEAPTSGSPGSAASPEVTPTSTPNPNNVTQNKTTPTEPASPPTTPKPTSTPKSPPTSTPDPKPKNNTTPAKSGRPTKP PGPVWCDRRDPLARYGSRVQIRCRFRNSTRMEFRLQIWRYSMGPSPPIAPAPDLEEVLTNITAPPGGLLVYDSAPNLTDPHVLWAEGAGPGADPPLYSVTGPLPTQRLIIGEVTPATQGMYYLAWGRMDSPHEYGTWVRVRMFRPPSLTLQPHAVMEGQPFKATCTAAAYYPRNPVEFVWFEDDHQVFNPGQIDTQTHEHPDGFTTVSTVTSEAVGGQVPPRTFTCQMTWHRDSVTFSRRNATGLALVLPRPTITMEFGVRIVVCTAGCVPEGVTFAWFLGDDPSPAAKSAVTAQESCDHPGLATVRSTLPISYDYSEYICRLTGYPAGIPVLEHHGSHQPPPRDPTERQVIEAIEWVGIGIGVLAAGVLVVTAIVYVVRTSQSRQRHRR

### 3.2. Antigenicity Prediction and Physicochemical Property Analysis of the Protein Sequences

From the selected nine proteins, only the antigenic proteins were selected for vaccine construction. VaxiJen v2.0 (http://www.ddg-pharmfac.net/vaxijen/VaxiJen/VaxiJen.htm) server was used for antigenicity determination. Among the nine protein, envelope glycoprotein E, envelope glycoprotein B and envelope glycoprotein D were determined as possible antigenic proteins by the server (**Table 01**). The physicochemical property analysis was conducted on these three proteins.

**Table 01.**
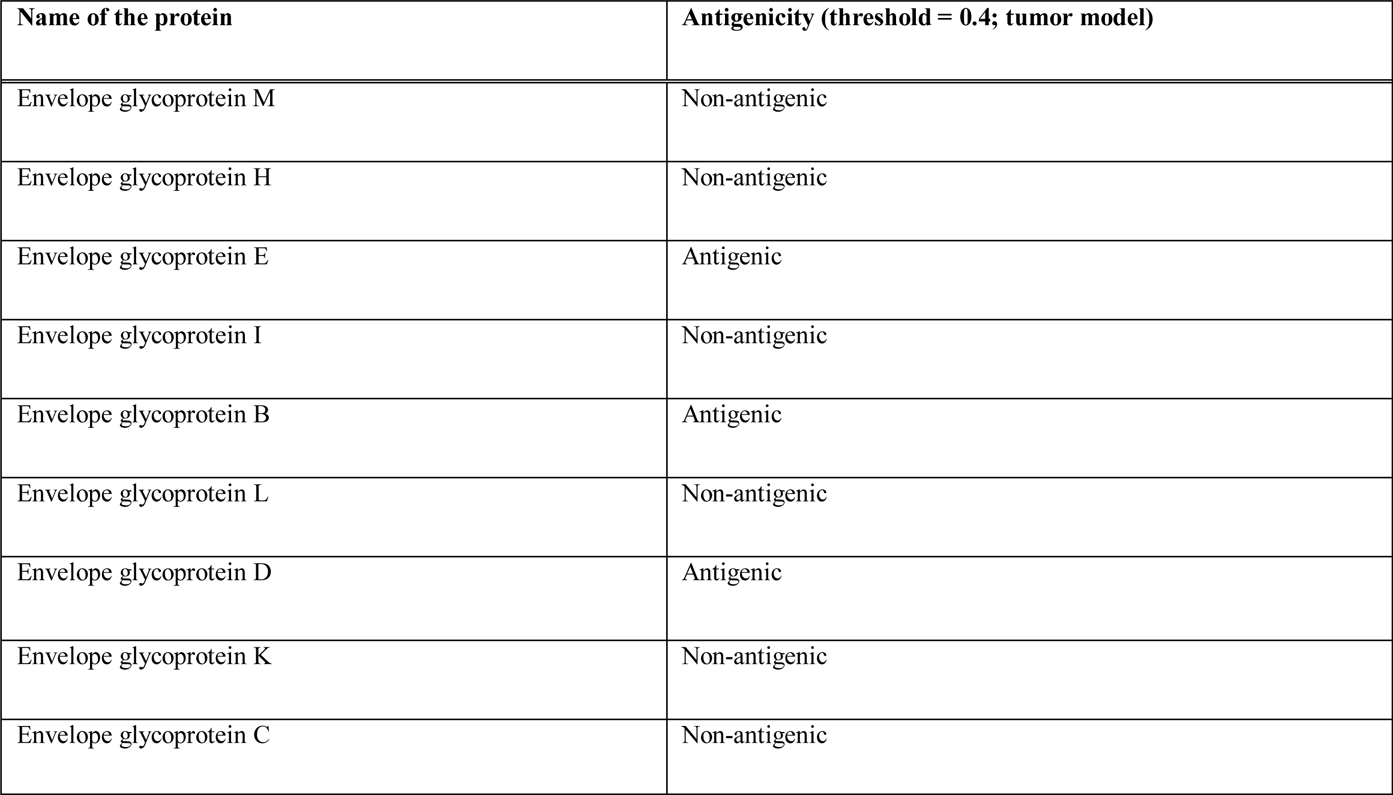
The antigenicity determination of the nine selected proteins.

The physicochemical study revealed that envelope glycoprotein B had the highest extinction co-efficient of 105255 M^-1^ cm^-1^ and lowest GRAVY value of −0.403. However, all the three proteins showed instability and all of them had half-lives of 30 hours in the mammalian cell culture system. The results of the physicochemical property analysis are listed in **Table 02**.

**Table 02.**
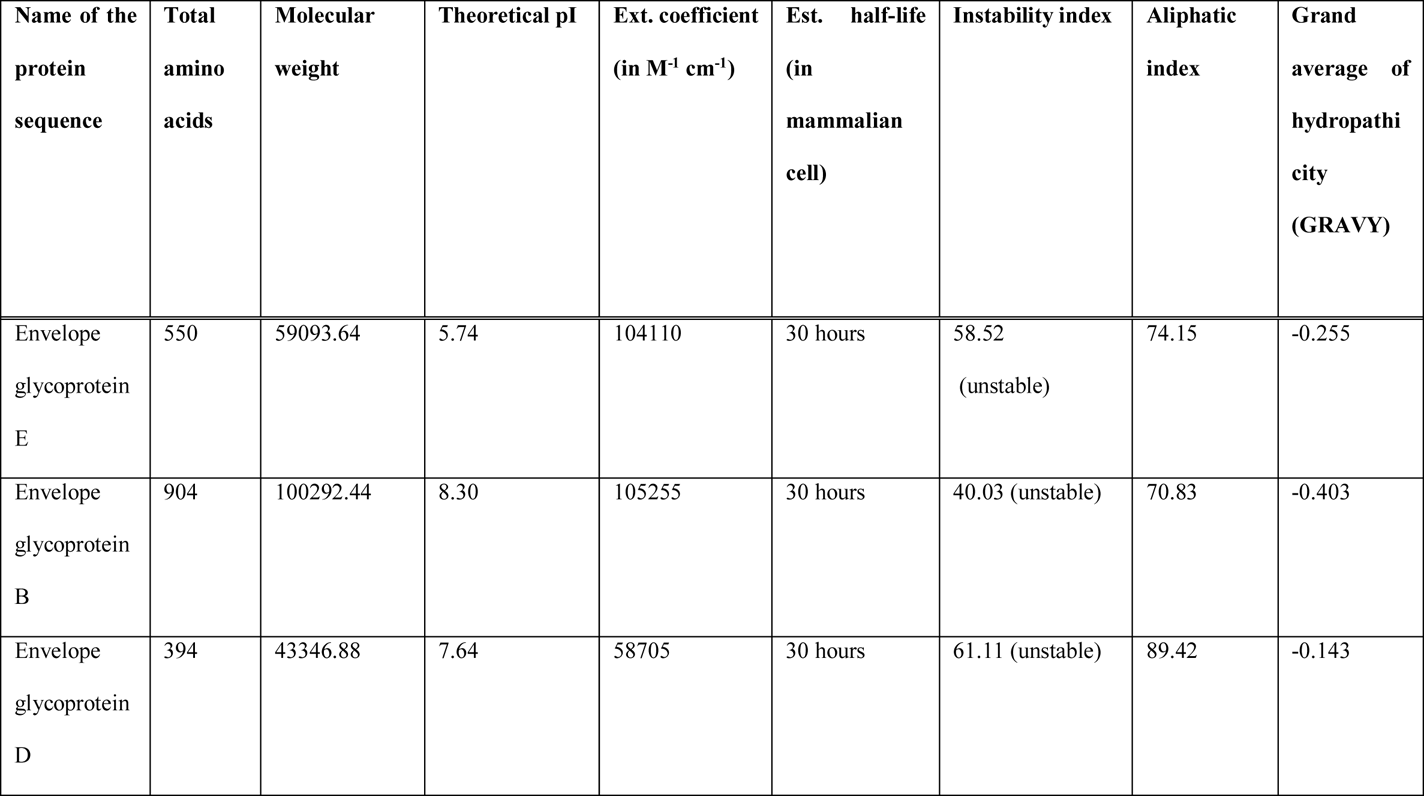
The antigenicity and physicochemical property analysis of the selected viral proteins.

### 3.3. T-cell and B-cell Epitope Prediction and Topology Determination of the Epitopes

The T-cell epitopes of MHC class-I of the three proteins were determined by NetMHCpan EL 4.0 prediction method of the IEDB (https://www.iedb.org/) server keeping the sequence length 09. The server yielded over 100 such epitopes. However, based on analyzing the best antigenicity scores (AS) and percentile scores for each epitope, the top ten potential epitopes were selected randomly for antigenicity, allergenicity, toxicity and conservancy tests. The server ranks the predicted epitopes based on the ascending order of percentile scores. The lower the percentile, the better the binding affinity. The T-cell epitopes of MHC class-II (HLA DRB1*04-01 allele) of the proteins were also determined by IEDB (https://www.iedb.org/) server, where the Sturniolo prediction method was used. Among the hundreds of epitopes generated by the server, best ten epitopes were selected for further analysis. Moreover, the B-cell epitopes of the proteins were selected using Bipipered linear epitope prediction method of the IEDB (https://www.iedb.org/) server and epitopes were selected based on their higher lengths (**Figure 02**). The topology of the selected epitopes were determined by TMHMM v2.0 server (http://www.cbs.dtu.dk/services/TMHMM/). **Table 03** and **Table 04** list the potential T-cell epitopes of envelope protein E, **Table 05** and **Table 06** list the potential T-cell epitopes of envelope glycoprotein B, **Table 07** and **Table 08** list the potential T-cell epitopes of envelope protein D and **Table 09** list the potential B-cell epitopes with their respective topologies.

**Figure 02.**
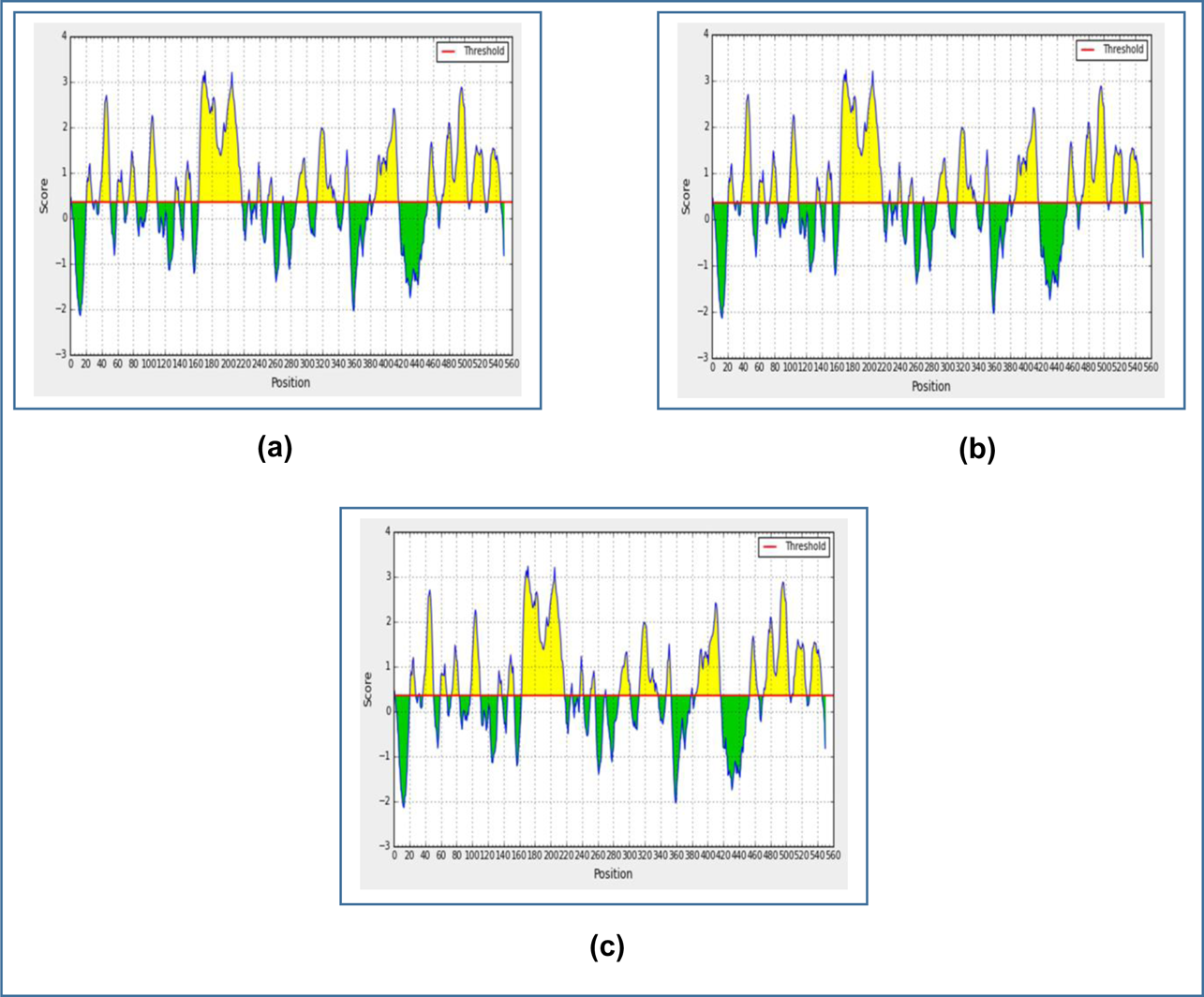
Figure showing the graphs of the B-cell epitope prediction of the three selected proteins of Human Herpes Simplex Virus-1, strain-17, using Bipipered linear epitope prediction method. Here, (a) is the graph of epitope prediction for envelope protein E, (b) is the graph of epitope prediction for envelope glycoprotein B and (c) is the graph of epitope prediction for envelope protein D.

**Table 03.**
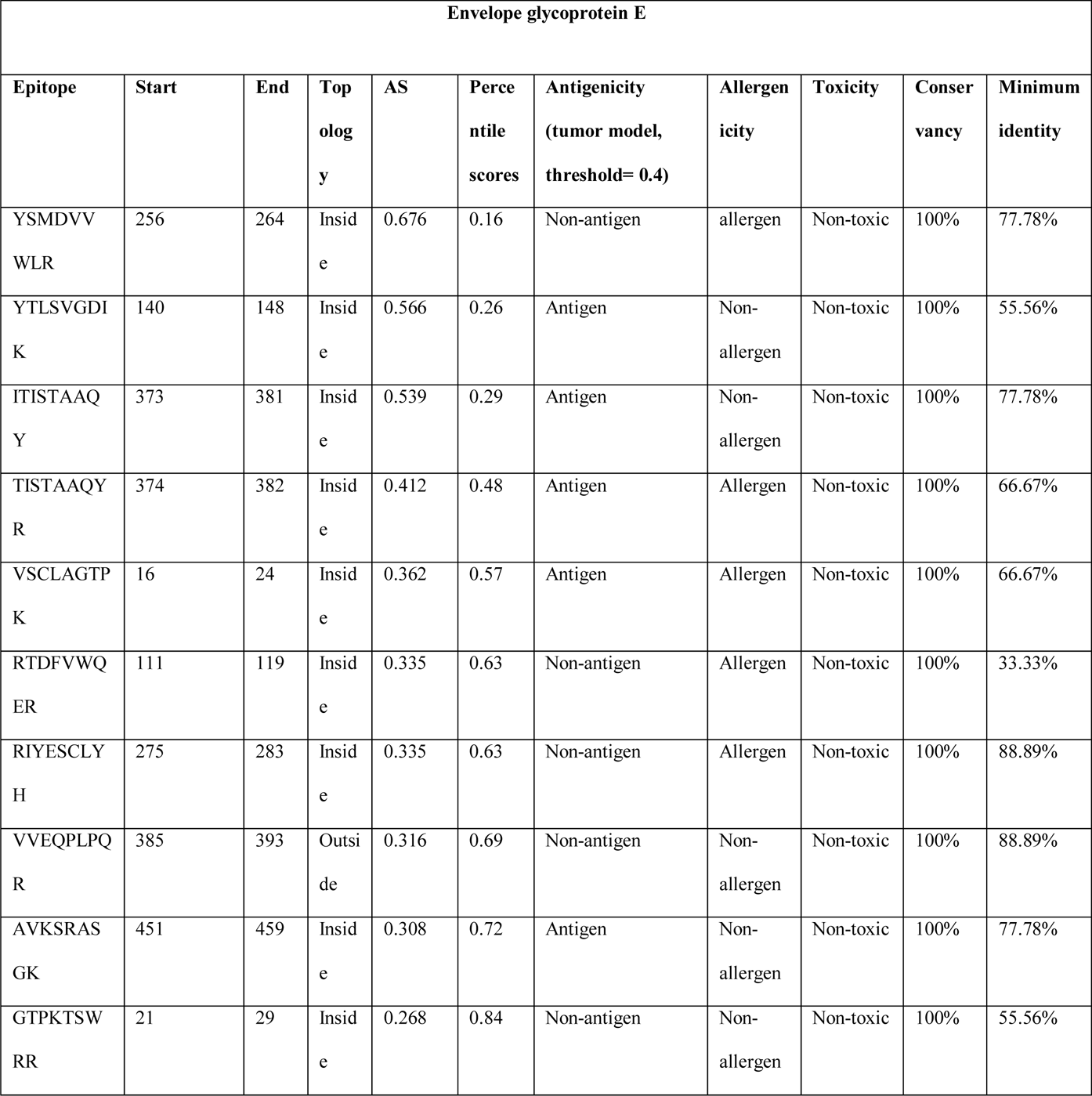
MHC class-I epitope prediction and topology, antigenicity, allergenicity, toxicity and conservancy analysis of the epitopes of envelope glycoprotein E.

**Table 04.**
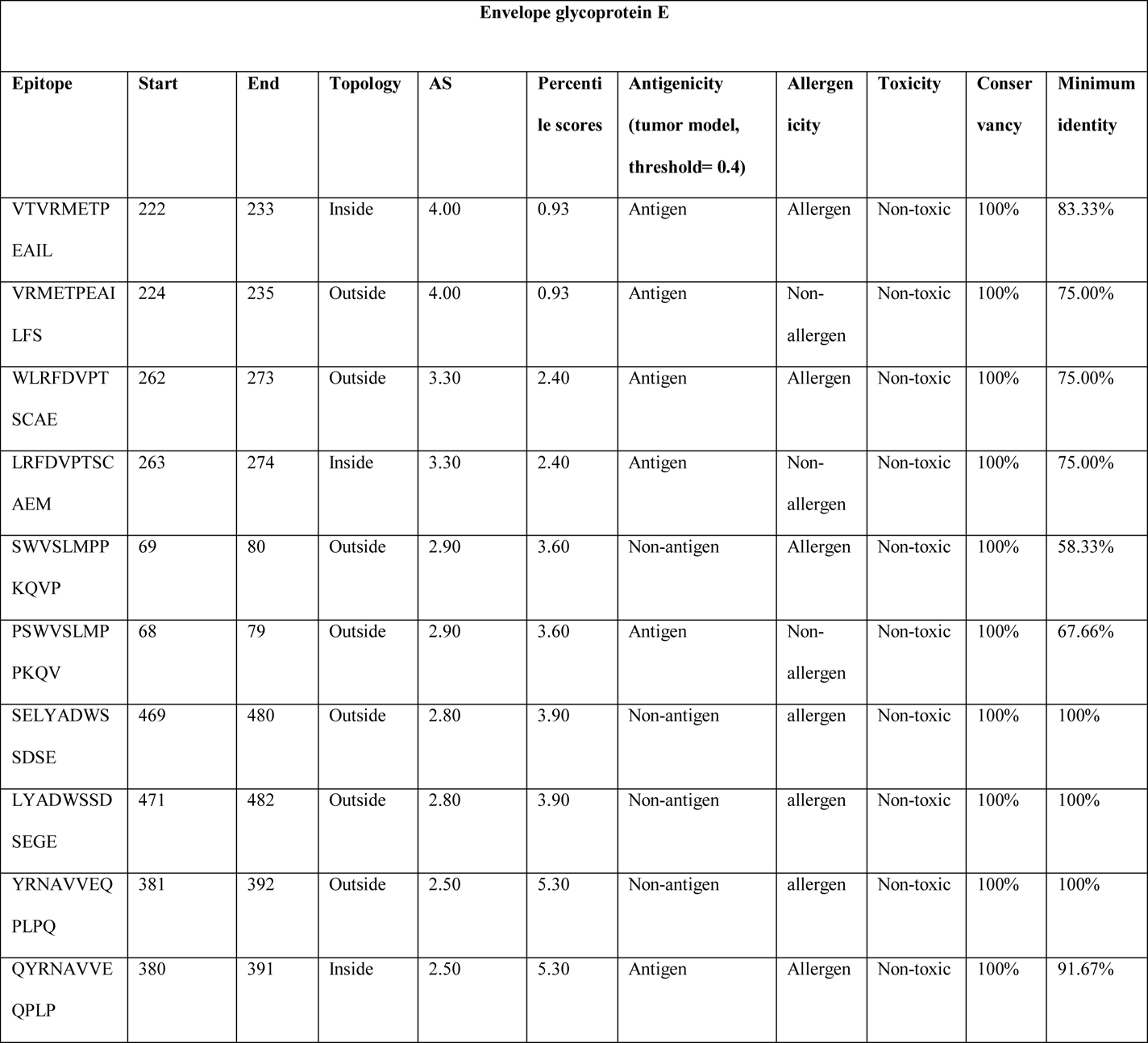
MHC class-II epitope prediction and topology, antigenicity, allergenicity, toxicity and conservancy analysis of the epitopes of envelope protein E.

**Table 05.**
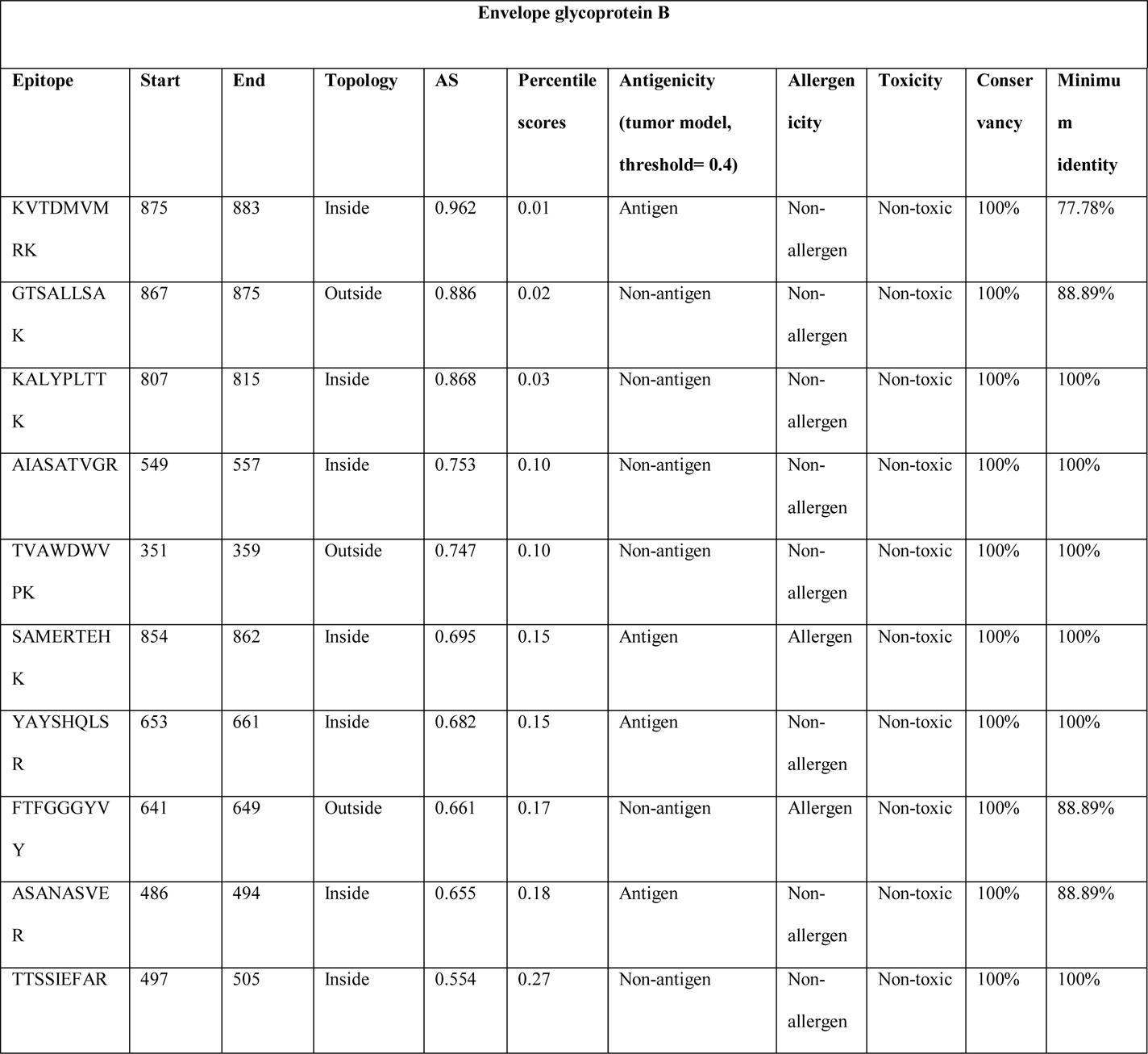
MHC class-I epitope prediction and topology, antigenicity, allergenicity, toxicity and conservancy analysis of the epitopes of envelope protein B.

**Table 06.**
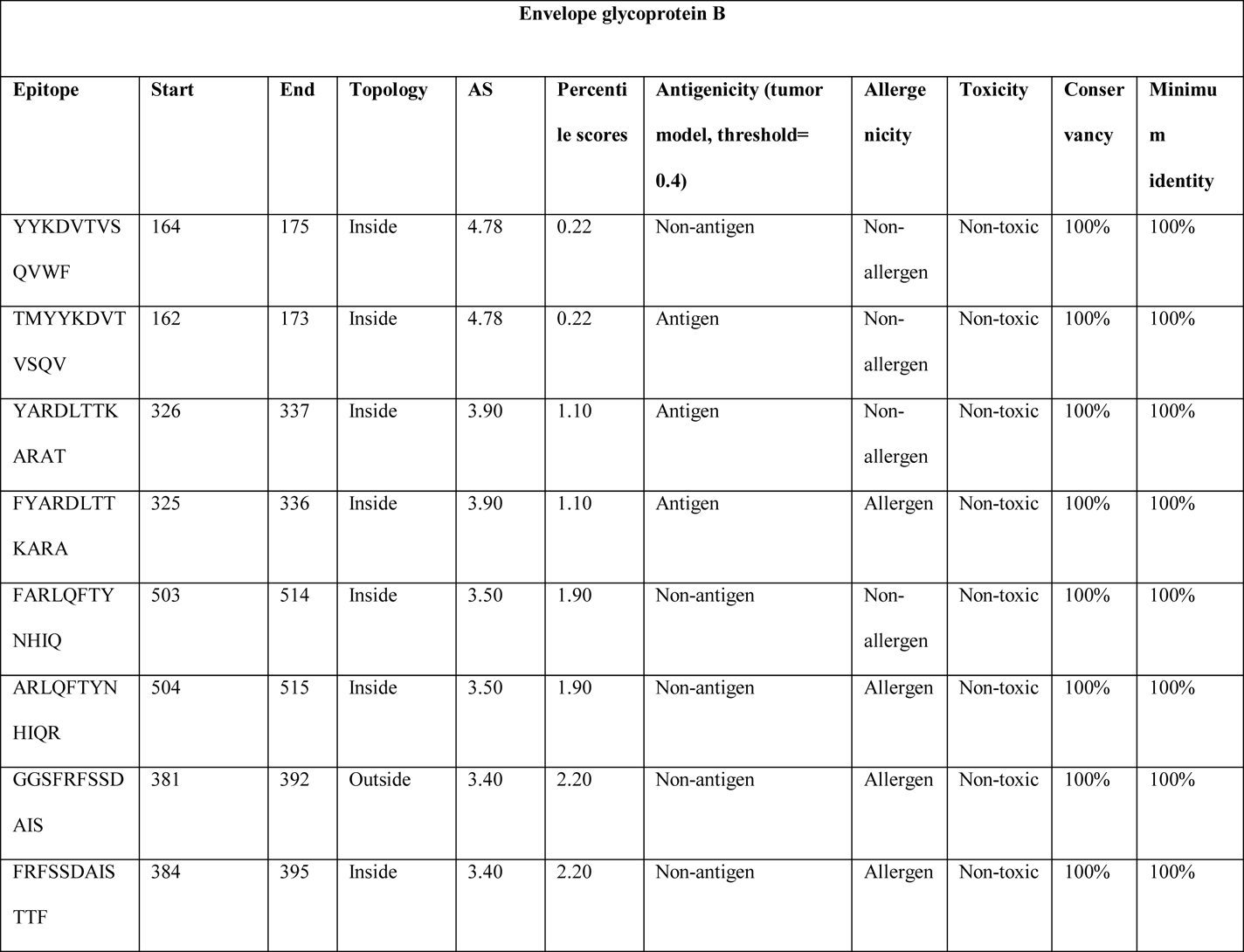
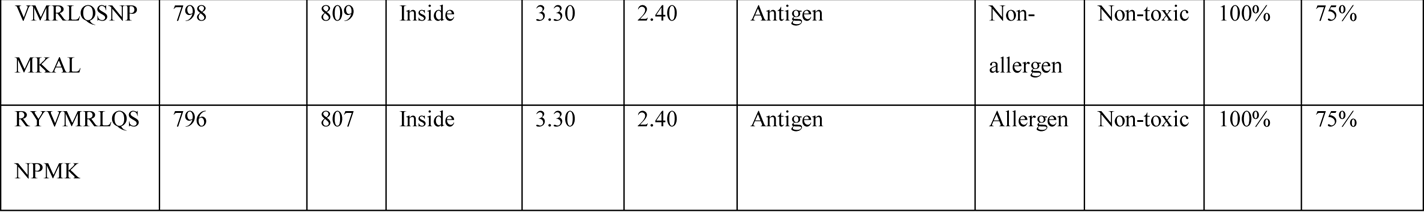
MHC class-II epitope prediction and topology, antigenicity, allergenicity, toxicity and conservancy analysis of the epitopes of envelope protein B.

**Table 07.**
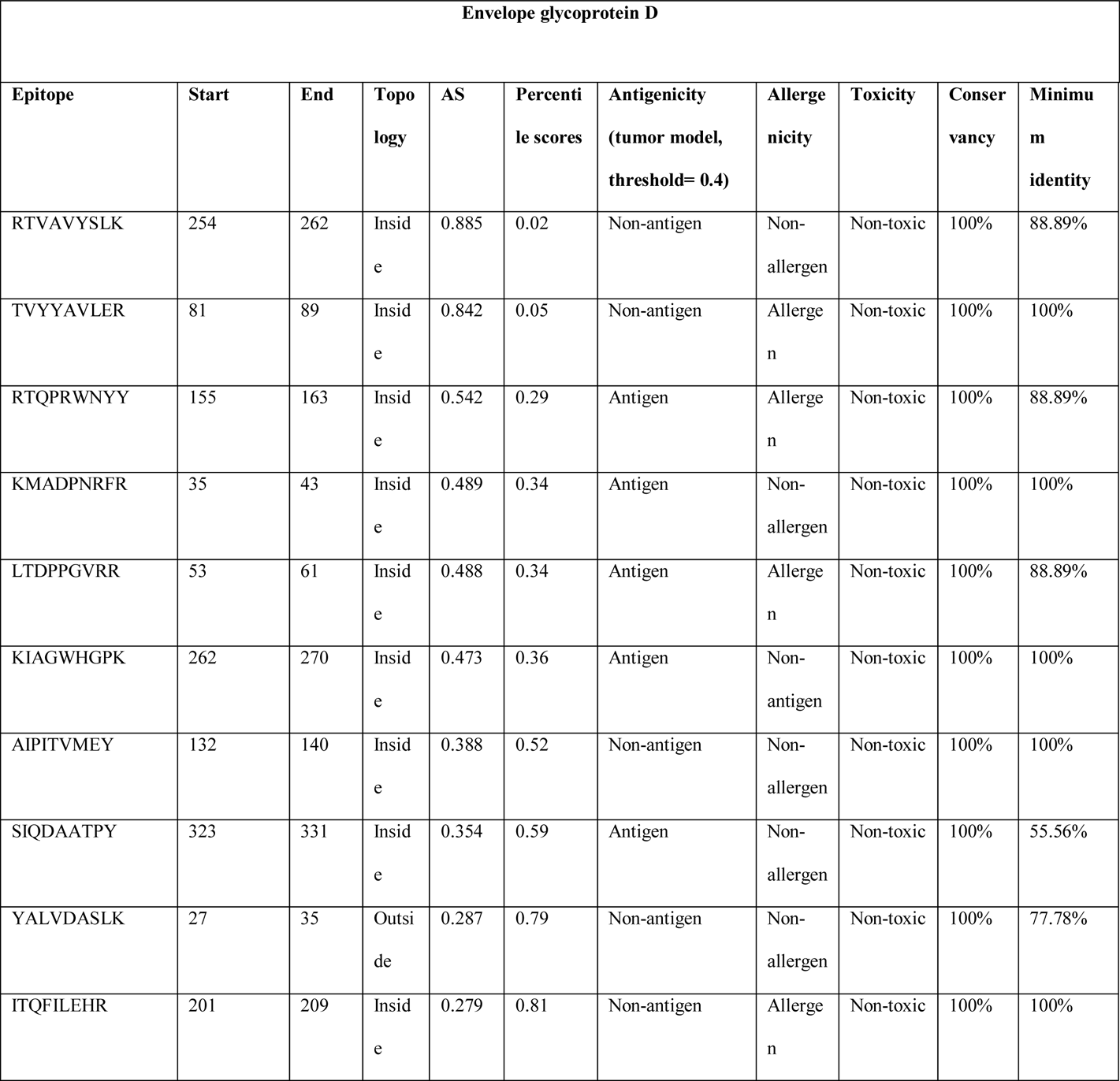
MHC class-I epitope prediction and topology, antigenicity, allergenicity, toxicity and conservancy analysis of the epitopes of envelope protein D.

**Table 08.**
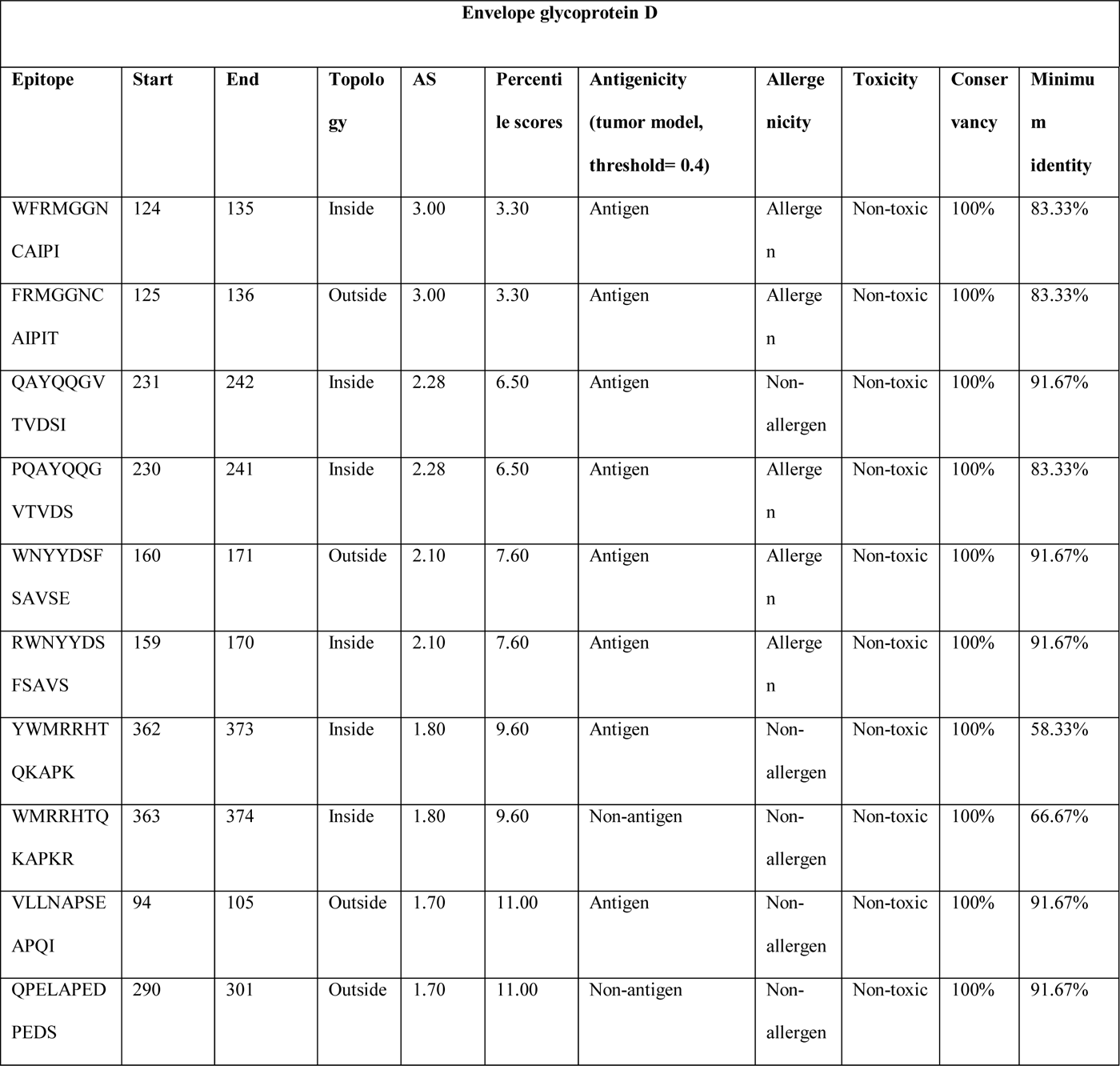
MHC class-II epitope prediction and topology, antigenicity, allergenicity, toxicity and conservancy analysis of the epitopes of envelope protein D.

**Table 09.**
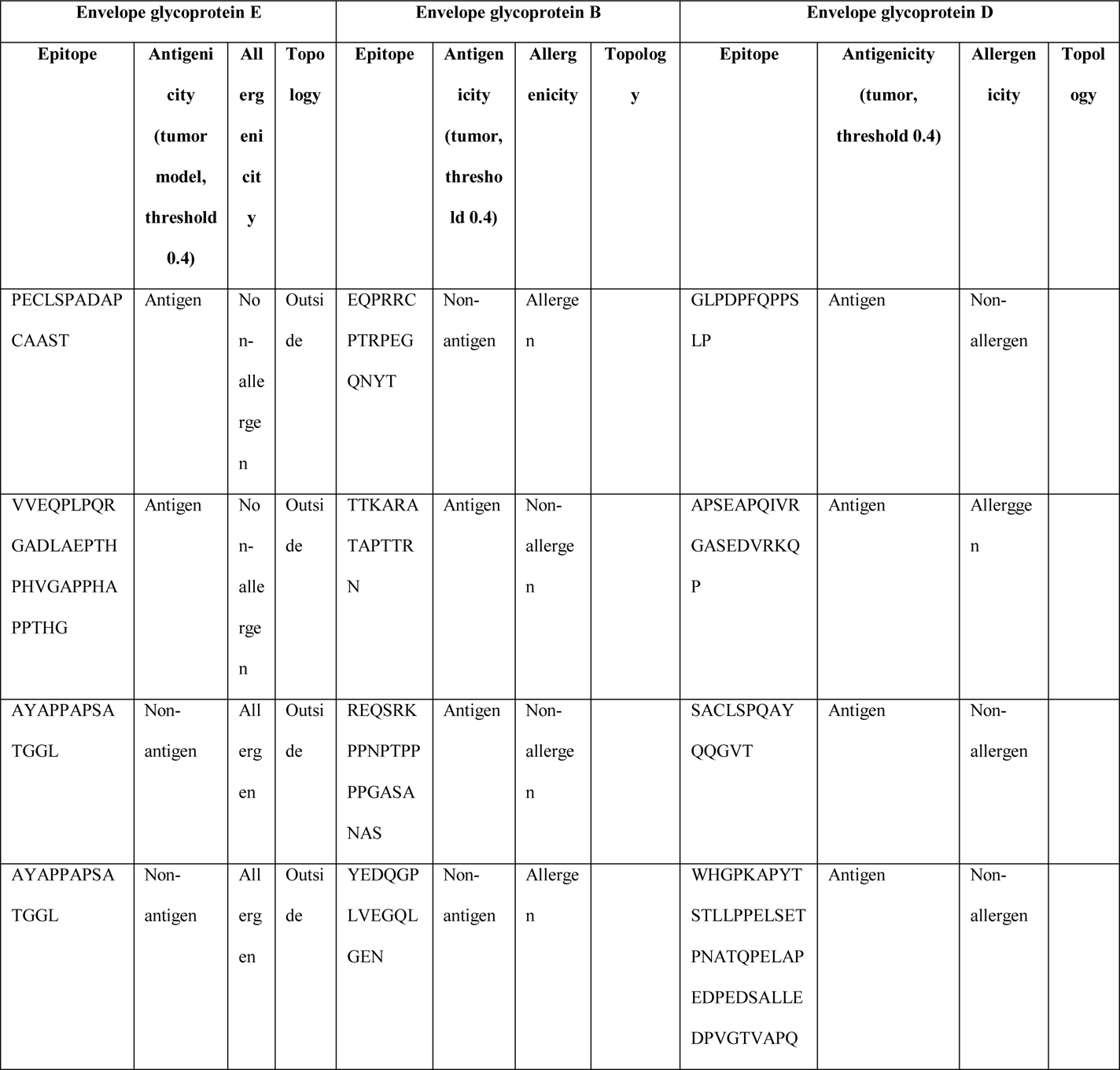
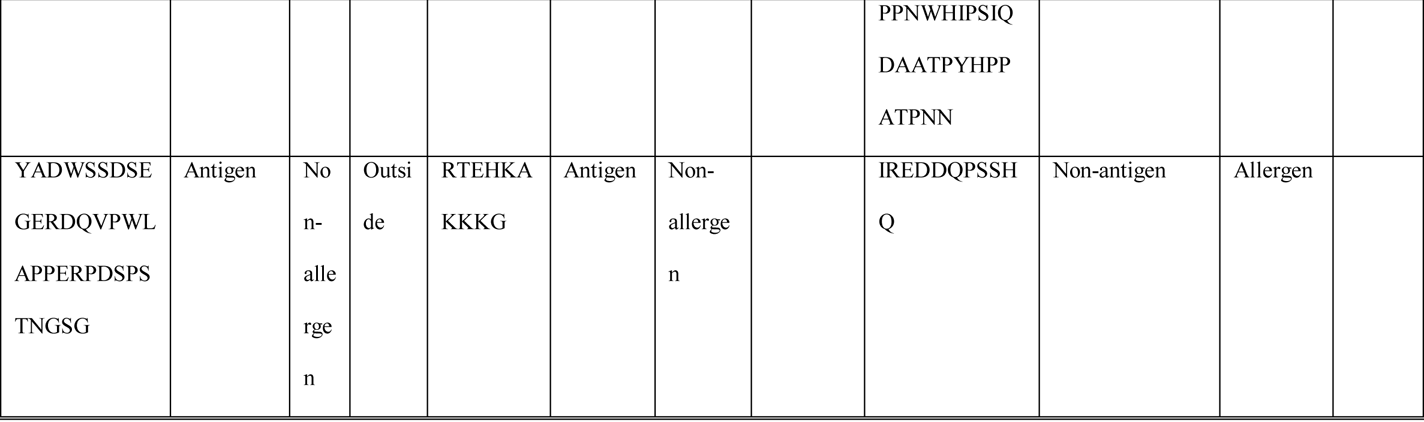
B-cell epitope prediction and antigenicity and allergenicity analysis of the epitopes of the three selected proteins.

### 3.4. Antigenicity, Allergenicity, Toxicity and Conservancy Analysis

In the antigenicity, allergenicity, toxicity and conservancy analysis, the T-cell epitopes that were found to be highly antigenic as well as non-allergenic, non-toxic, had minimum identity of over 50% and had conservancy of over 90% and the antigenic as well as non-allergenic B-cell epitopes were selected for further analysis and vaccine construction. Among the ten selected MHC class-I epitopes and ten selected MHC class-II epitopes of envelope glycoprotein E, total six epitopes (three epitopes from each of the category) were selected based on the mentioned criteria: YTLSVGDIK, ITISTAAQY, AVKSRASGK, VRMETPEAILFS, LRFDVPTSCAEM and PSWVSLMPPKQV. On the other hand, among the ten selected MHC class-I epitopes and ten selected MHC class-II epitopes of envelope glycoprotein B, total six epitopes (three epitopes from each of the category) were selected based on the mentioned criteria: KVTDMVMRK, YAYSHQLSR, ASANASVER, TMYYKDVTVSQV, YARDLTTKARAT and VMRLQSNPMKAL. Moreover, like these proteins, six epitopes that obeyed the mentioned criteria, were selected for further analysis from the envelope glycoprotein D epitopes: KMADPNRFR, KIAGWHGPK, SIQDAATPY, QAYQQGVTVDSI, YWMRRHTQKAPK and VLLNAPSEAPQI. For the selection of the B-cell epitopes, the highly antigenic and non-allergenic sequences were taken for vaccine construction. Three epitopes from each of the protein category were selected. For this reason, total nine epitopes B-cell epitopes were selected for vaccine construction, since they obeyed the selection criteria. (**Table 03**, **Table 04**, **Table 05**, **Table 06**, **Table 07**, **Table 08** and **Table 09**).

### 3.5. Cluster Analysis of the MHC Alleles

The cluster analysis of the possible MHC class-I and MHC class-II alleles that may interact with the predicted epitopes of the three selected proteins, were performed by online tool MHCcluster 2.0 (http://www.cbs.dtu.dk/services/MHCcluster/). The tool generates the relationship of the clusters of the alleles in phylogenetic manner. **Figure 03** illustrates the outcome of the experiment where the red zone indicates strong interaction and the yellow zone corresponds to weaker interaction.

**Figure 03.**
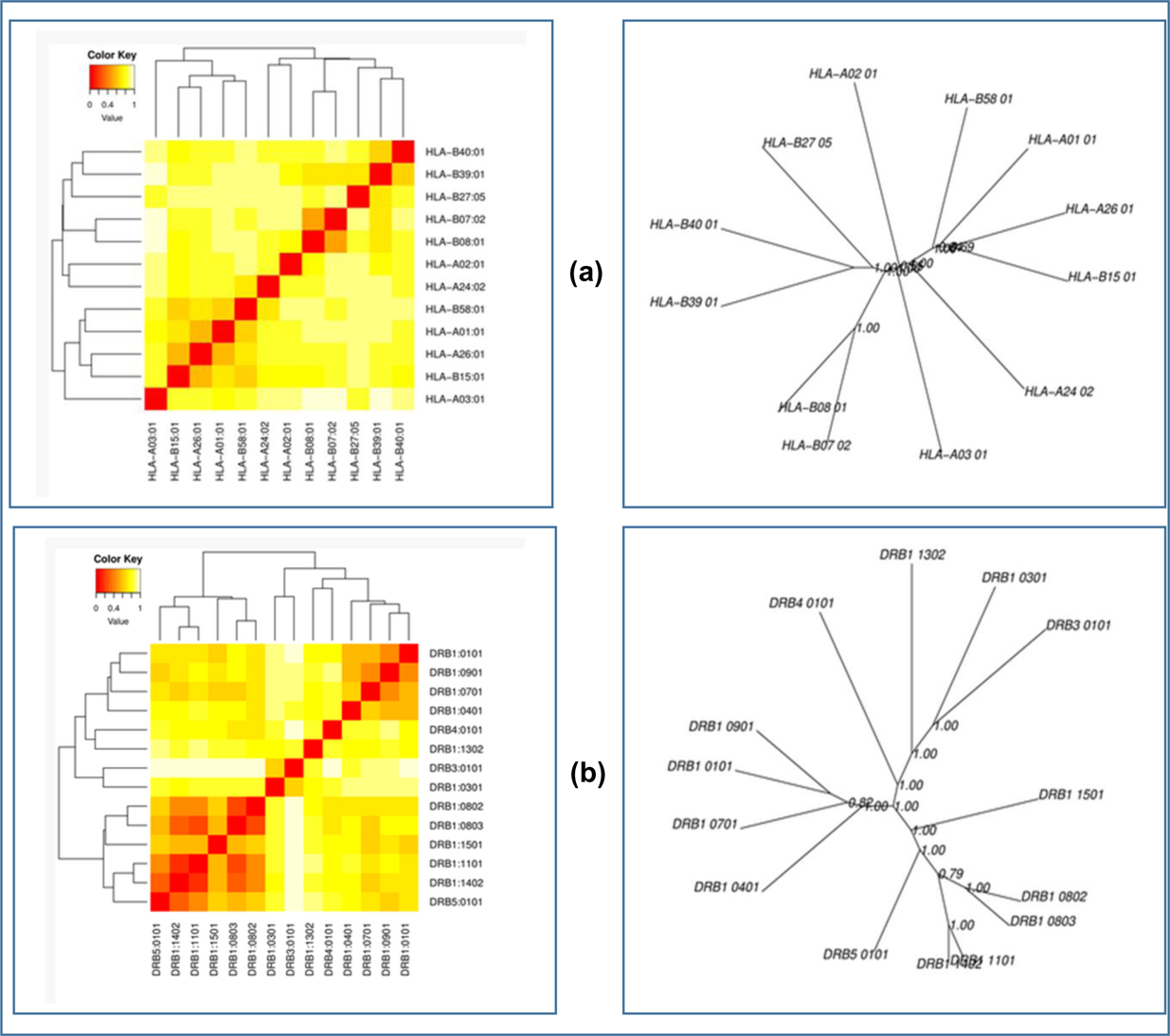
The results of the MHC cluster analysis. Here, (a) is the heat map (left) and the tree map (right) of MHC class-I cluster analysis, (b) is the heat map (left) and the tree map (right) of MHC class-II cluster analysis. The cluster analysis was carried out using onine server MHCcluster 2.0 (http://www.cbs.dtu.dk/services/MHCcluster/).

**Figure 04.**
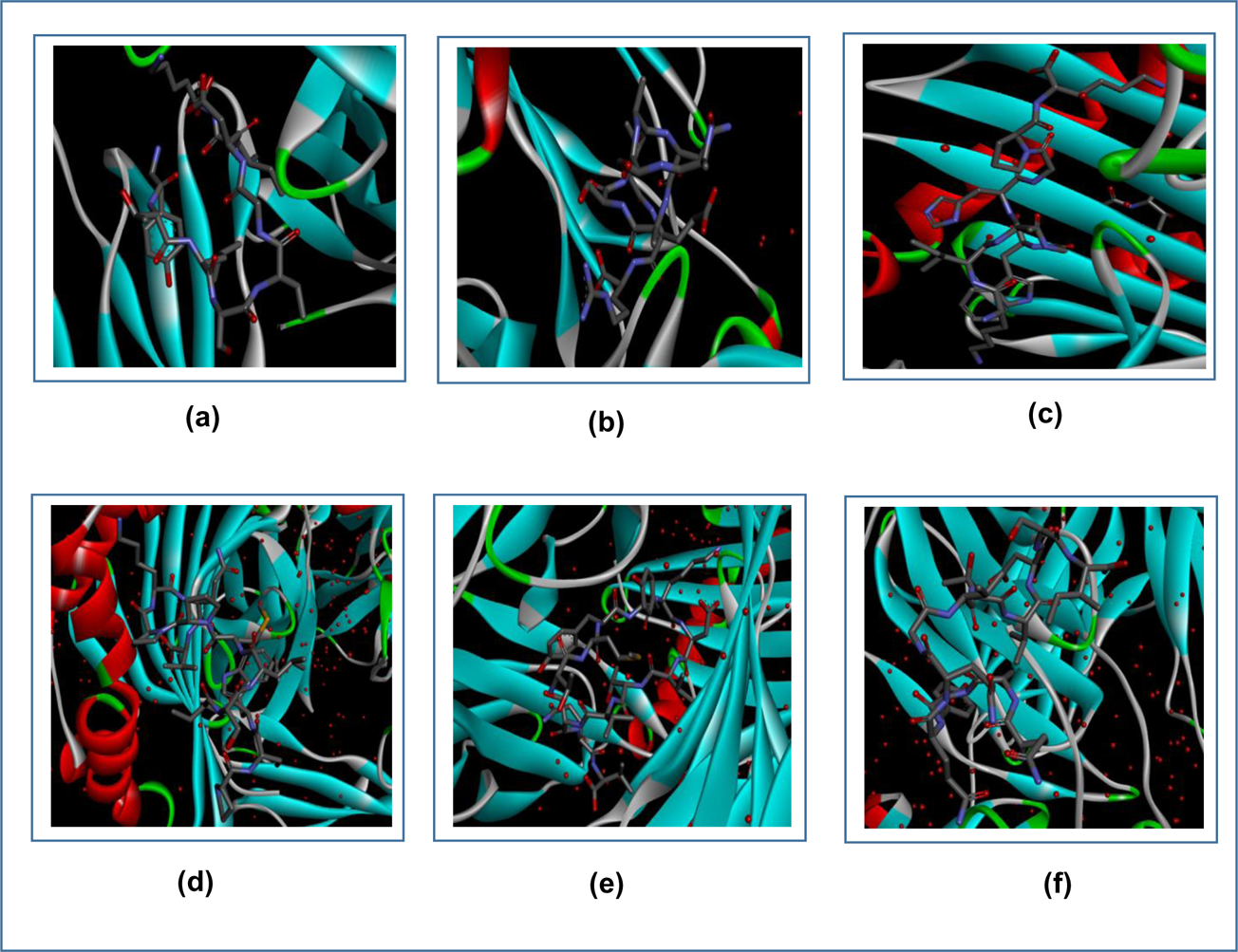
Figure showing the interactions between the best epitopes from the three proteins and their respective receptors. Here, (a) is the interaction between YTLSVGDIK and MHC class-I, (b) is the interaction between ASANASVER and MHC class-I, (c) is the interaction between KIAGWHGPK and MHC class-I molecule, (d) is the interaction between PSWVSLMPPKQV and MHC class-II, (e) is the interaction between TMYYKDVTVSQV and MHC class-II, (f) is the interaction between QAYQQGVTVDSI and MHC class-II molecule. The interactions were visualized by Discovery Studio Visualizer.

### 3.6. Generation of the 3D Structures of the Epitopes and Peptide-Protein Docking

The 3D structures of the selected T-cell epitopes were generated by the PEP-FOLD3 server. The 3D structures were generated for peptide-protein docking. The docking was carried out to find out, whether all the epitopes had the capability to bind with the MHC class-I and MHC class-II molecules. All the MHC class-I epitopes were docked against the HLA-A*11-01 allele (PDB ID: 5WJL) and MHC class-II HLA DRB1*04-01 (PDB ID: 5JLZ). The docking was performed using PatchDock server and then the results were refined by FireDock online server. Among the MHC class-I epitopes of envelope glycoprotein E, YTLSVGDIK showed the best result with the lowest global energy of −41.93. Among the MHC class-I epitopes of envelope glycoprotein B, ASANASVER generated the lowest and best global energy score of −36.46. KIAGWHGPK generated the best global energy score of −41.47 of the MHC class-I epitopes of envelope glycoprotein D. Among the MHC class-II epitopes of envelope glycoprotein E, PSWVSLMPPKQV generated the best global energy score of −7.76. TMYYKDVTVSQV generated the lowest global energy of −54.94 and QAYQQGVTVDSI generated the lowest global energy of −4.25, among the MHC class-II epitopes of envelope glycoprotein B and envelope glycoprotein D, respectively (**Table 10**).

**Table 10.**
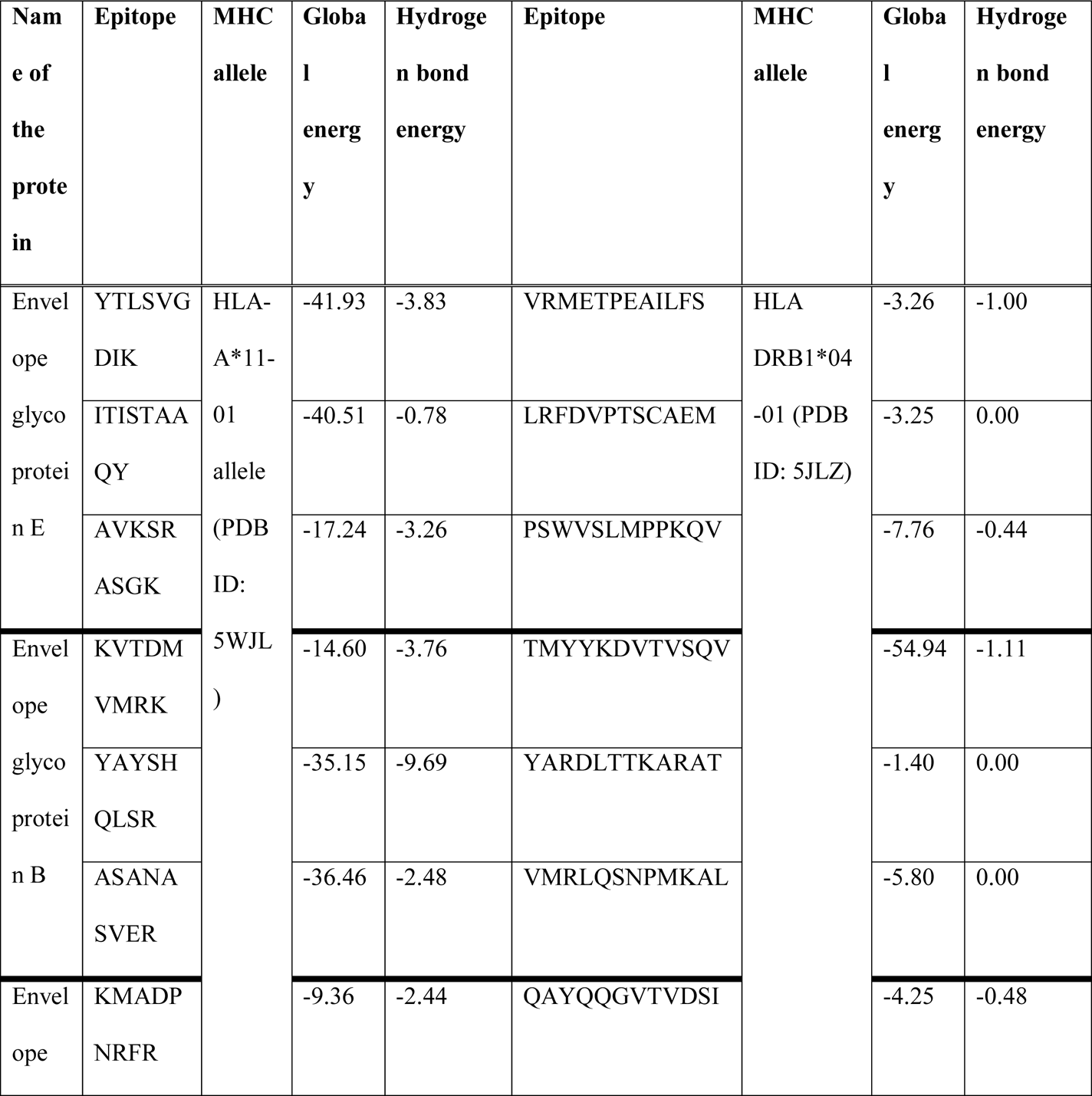
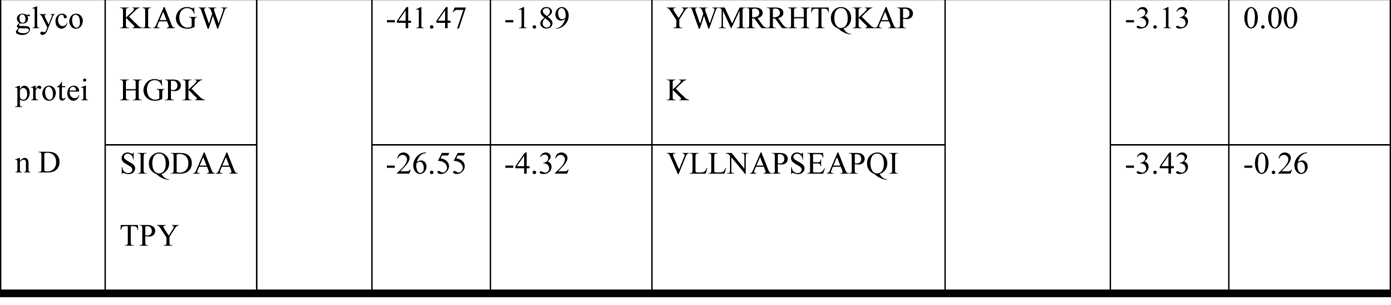
Results of molecular docking analysis of the selected epitopes.

### 3.7. Vaccine Construction

After successful docking, three vaccines were constructed, that could be used effectively to fight against HSV-1, strain-17. For vaccine constructions, three different adjuvants were used to construct three different vaccine for each of the viruses: beta defensin, L7/L12 ribosomal protein and HABA protein, were used. These three vaccines differ from each other only in their adjuvant sequences. PADRE sequence was also used for vaccine construction. Three different vaccine constructs differed from each other only in their adjuvant sequences. During vaccine construction, EAAAK, GGGS, GPGPG and KK linkers were used at their required positions. Each vaccine construct was ended by an additional GGGS linker. The newly constructed vaccines were designated as: HV-1, HV-2 and HV-3 (**Table 11**).

**Table 11.**
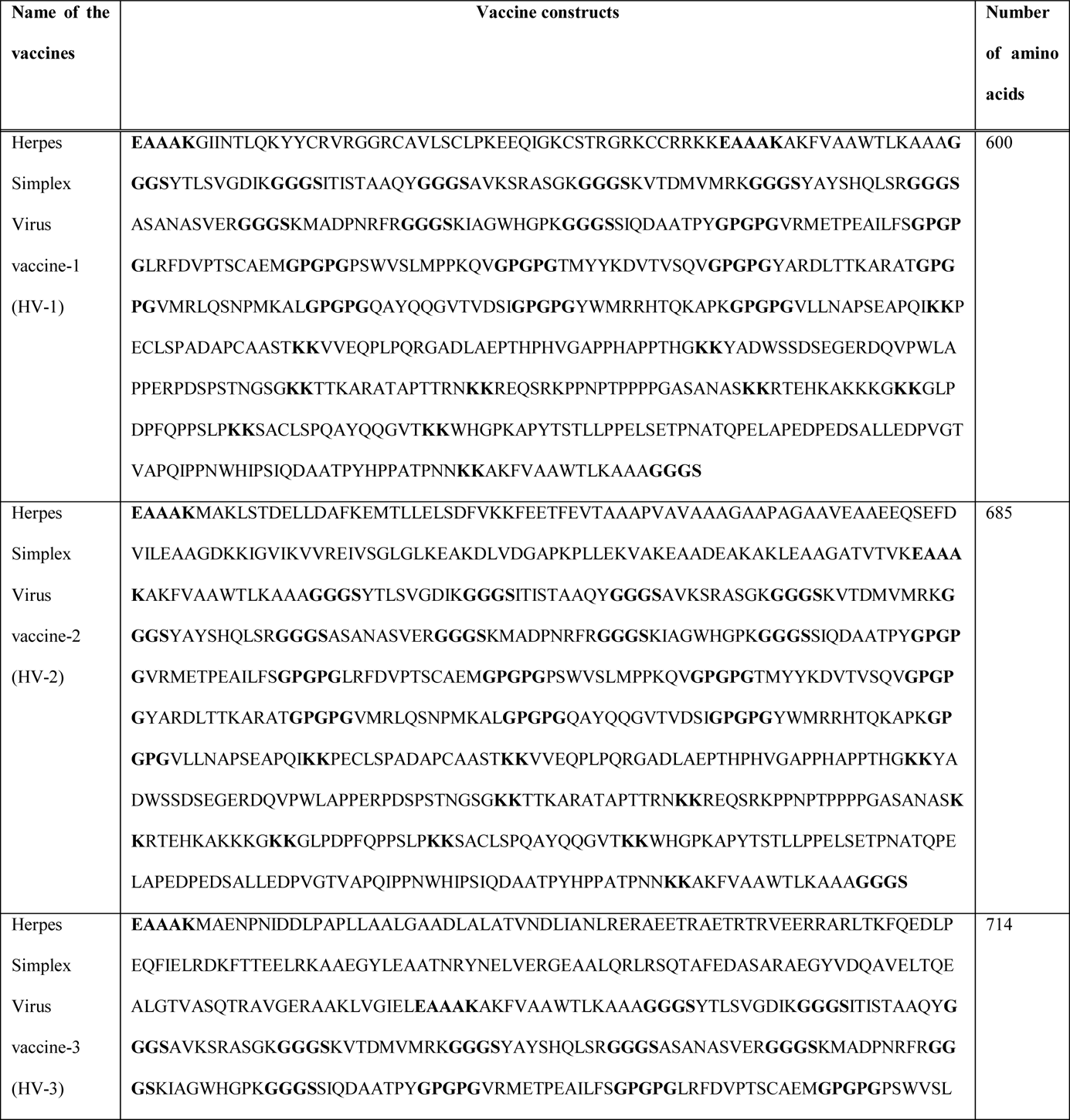
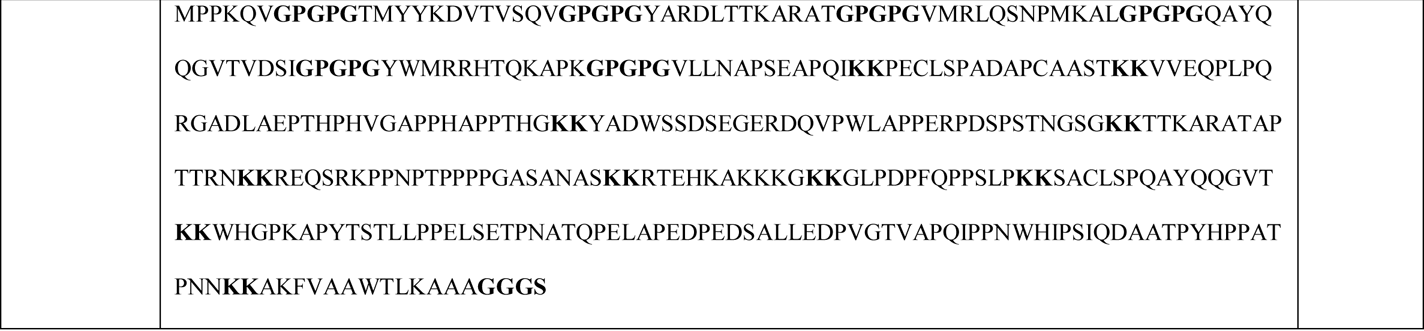
List of the vaccines constructed against Herpes Simplex Virus-1, strain-17.

### 3.8. Antigenicity, Allergenicity and Physicochemical Property Analysis of the Vaccine Constructs

The three vaccines were found to be potent antigen as well as non-allergen. Since they are found to be non-allergenic, they are safe to use. In the physicochemical property analysis, the number of amino acids, molecular weight, extinction coefficient (in M-1 cm-1), theoretical pI, half-life, aliphatic index and GRAVY were determined. All the vaccines quite similar theoretical pI and ext. coefficient (HV-3 had the lowest value of 74566.83 M-1 cm-1). All of the vaccine constructs had the similar half-life of 1 hour in the mammalian cells. HV-2 had the highest GRAVY value of −0.544. The antigenicity, allergenicity and physicochemical property analysis of the three vaccine constructs are listed in **Table 12**.

**Table 12.**
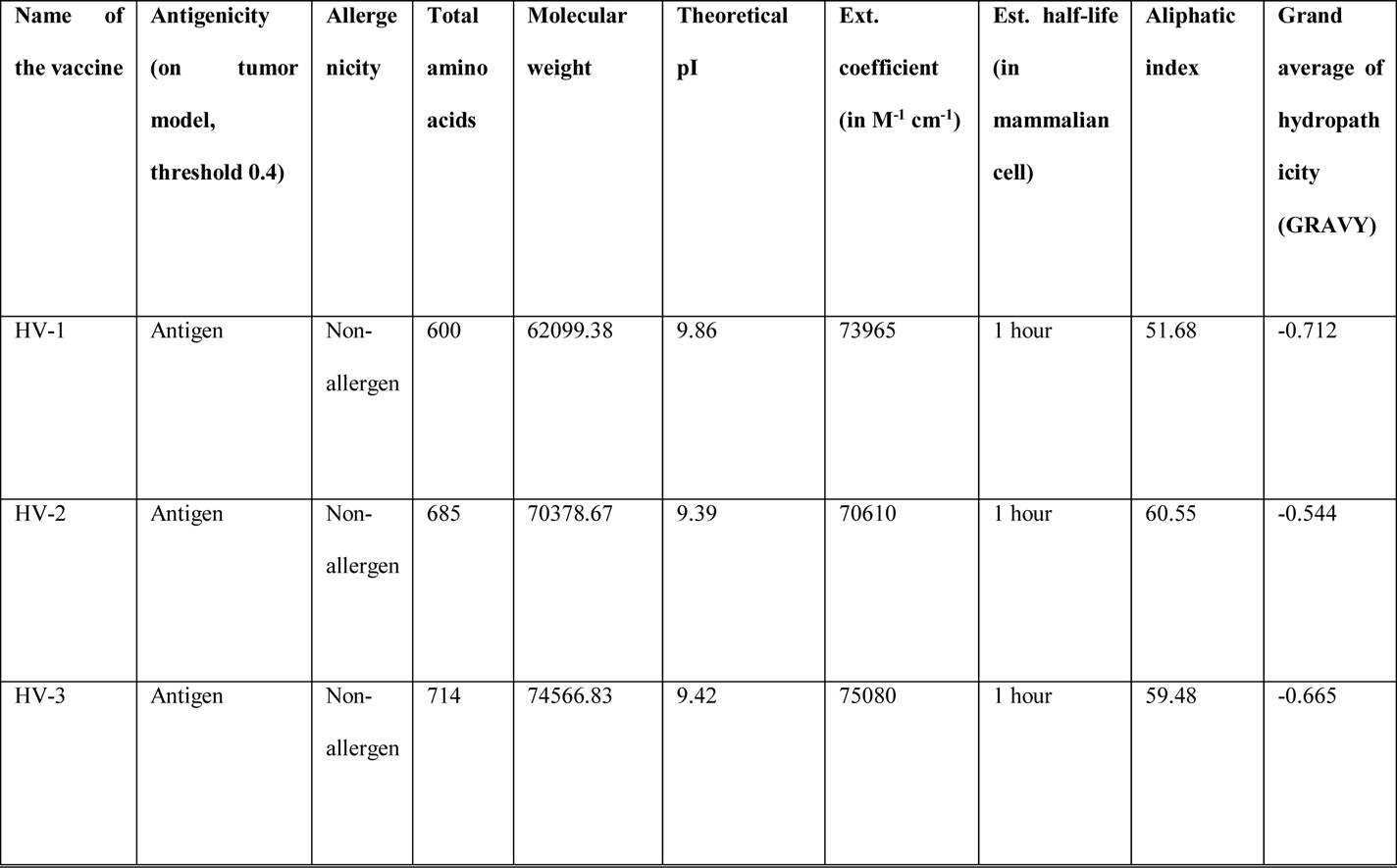
Antigenicity, allergenicity and physicochemical property analysis of the three vaccine constructs.

### 3.9. Secondary and Tertiary Structure Prediction of the Vaccine Constructs

The secondary structures of the three vaccine constructs were generated by the online tools, PRISPRED (http://bioinf.cs.ucl.ac.uk/psipred/) and NetTurnP v1.0 (http://www.cbs.dtu.dk/services/NetTurnP/). From the secondary structure analysis, it was analyzed that, the HV-1 had the highest percentage of the amino acids (74.1%) in the coil formation as well as the highest percentage of amino acids (10.8%) in the beta-strand formation. However, HV-3 had the highest percentage of 30.1% of amino acids in the alpha-helix formation (**Figure 05** and **Table 13**).

**Figure 05.**
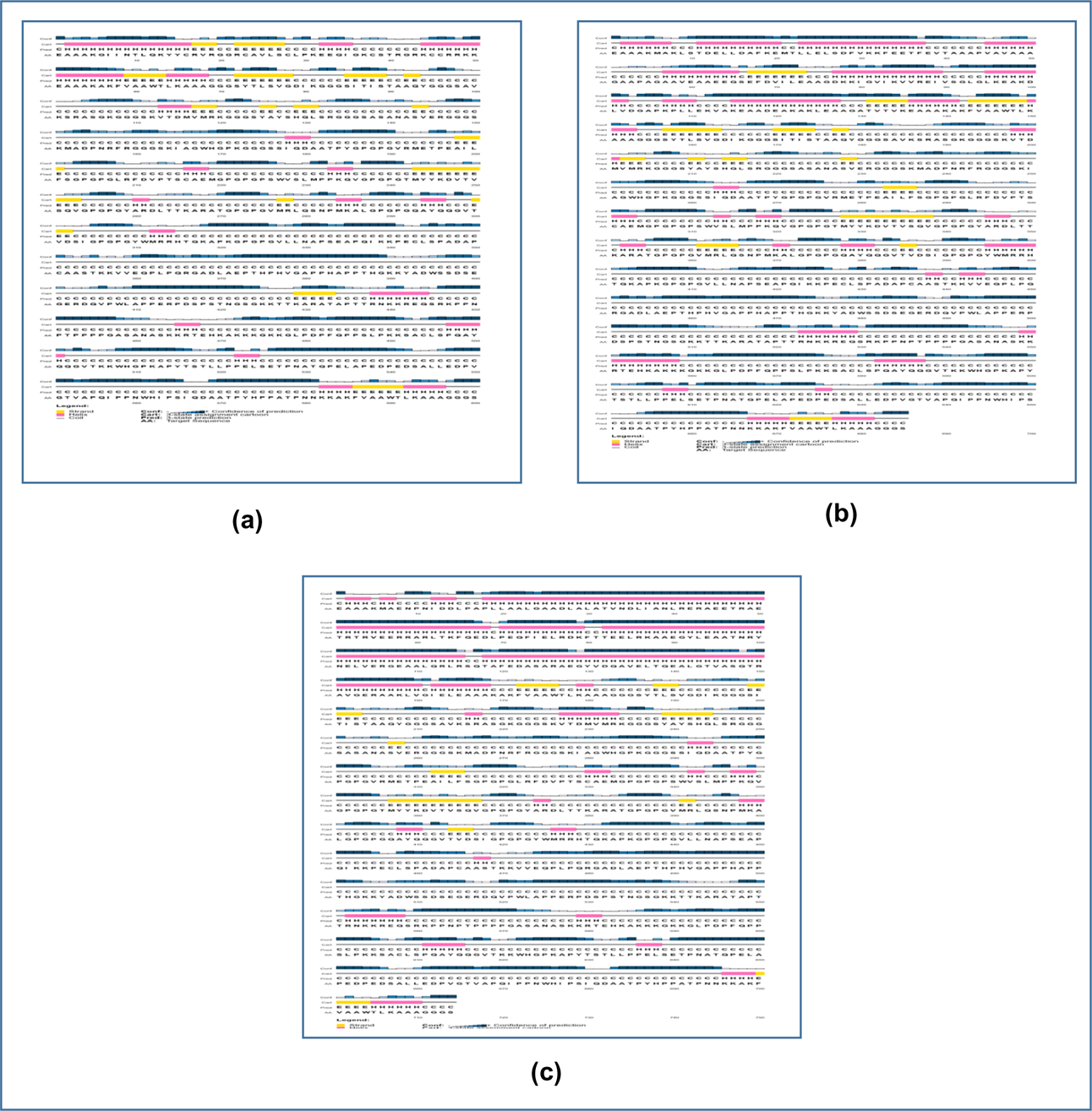
The results of the secondary structure prediction of the constructed Herpes vaccines. Here, (a) is HV-1, (b) is HV-2, (c) is HV-3. The secondary structures were predicted using online server PRISPRED (http://bioinf.cs.ucl.ac.uk/psipred/).

**Table 13.**
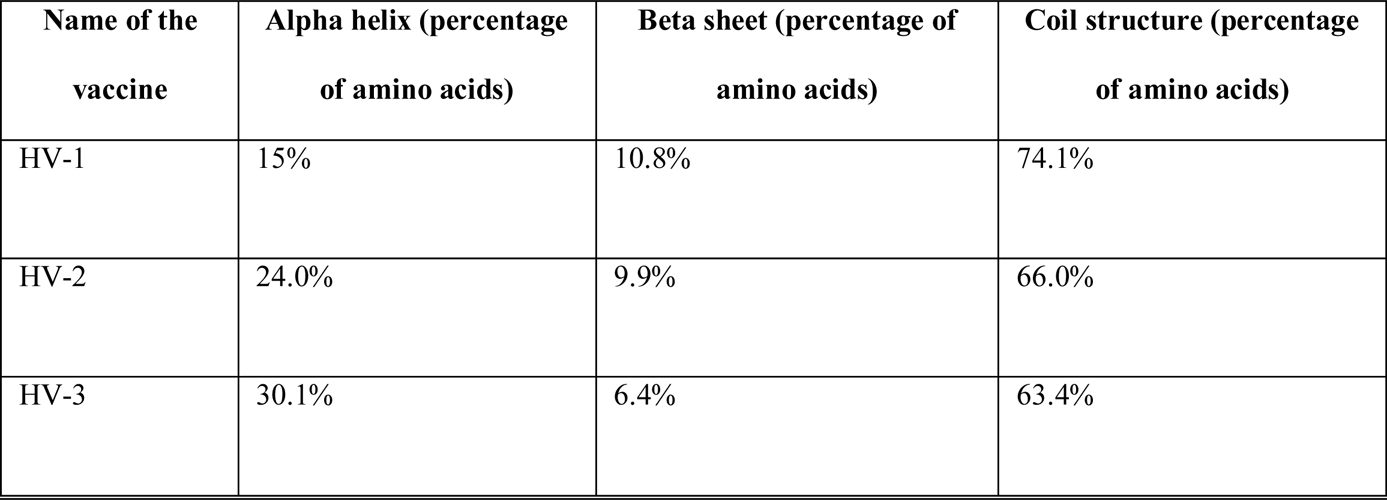
Results of the secondary structure analysis of the vaccine constructs.

The 3D structures of the vaccine constructs were predicted by the online server RaptorX (http://raptorx.uchicago.edu/). All the three vaccines had 3 domains and HV-2 had the lowest p-value of 8.91e-05. The homology modeling of the three dengue vaccine constructs were carried out using 1KJ6A (for HV-1), 1DD3A (for HV-2) and 4TQLA (for HV-3) as templates from protein data bank (https://www.rcsb.org/). The results of the 3D structure analysis are listed in **Table 14** and illustrated in **Figure 06**.

**Figure 06.**
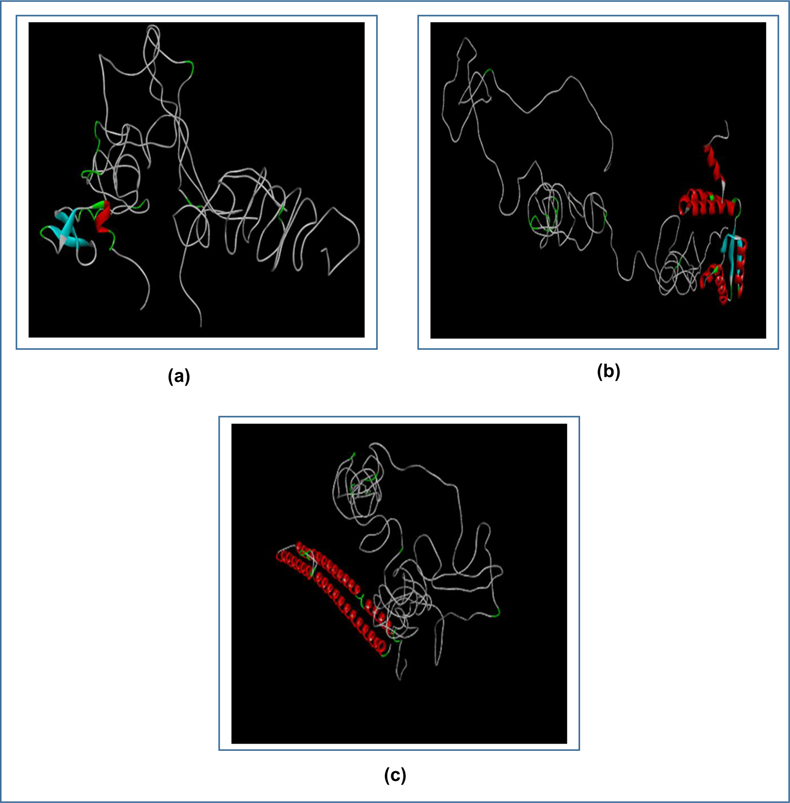
The tertiary structures of the three Herpes vaccines. Here, (a) is the HV-1, (b) is the HV-2 and (c) is the HV-3. The tertiary structures were predicted using the online server tool RaptorX (http://raptorx.uchicago.edu/) and visualized by Discovery Studio Visualizer.

**Table 14.**
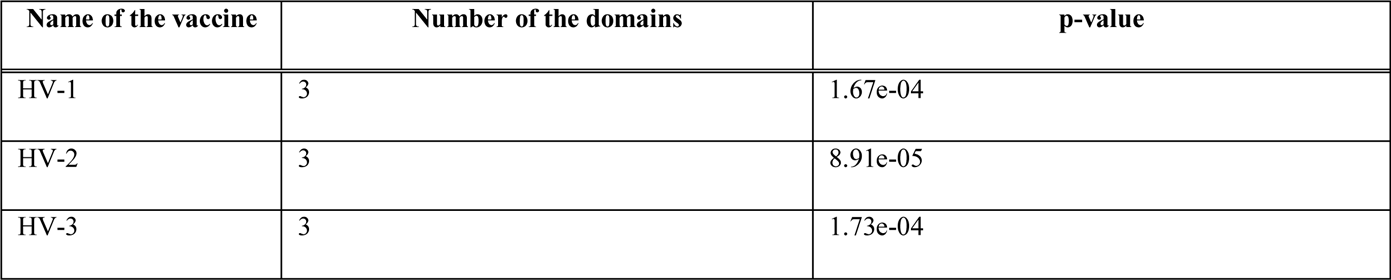
Results of the tertiary structure analysis of the vaccine constructs.

### 3.10. Protein 3D Structure Refinement and Validation

The protein structures generated by the RaptorX server (http://raptorx.uchicago.edu/) were refined using 3Drefine (http://sysbio.rnet.missouri.edu/3Drefine/) and then the refined structures were analyzed by Ramachandran plot generated by PROCHECK server (https://servicesn.mbi.ucla.edu/PROCHECK/). The analysis showed that HV-1 vaccine had 63.9% of the amino acids in the most favored region, 30.6% of the amino acids in the additional allowed regions, 3.7% of the amino acids in the generously allowed regions and 1.8% of the amino acids in the disallowed regions. The HV-2 vaccine had 70.4% of the amino acids in the most favored regions, 26.2% of the amino acids in the additional allowed regions, 2.9% of the amino acids in the generously allowed regions and 0.5% of the amino acids in the disallowed regions. The HV-3 vaccine had 74.7% of the amino acids in the most favored regions, 23.7% of the amino acids in the additional allowed regions, 1.1% of the amino acids in the generously allowed regions and 0.5% of the amino acids in the disallowed regions (**Figure 07**).

**Figure 07.**
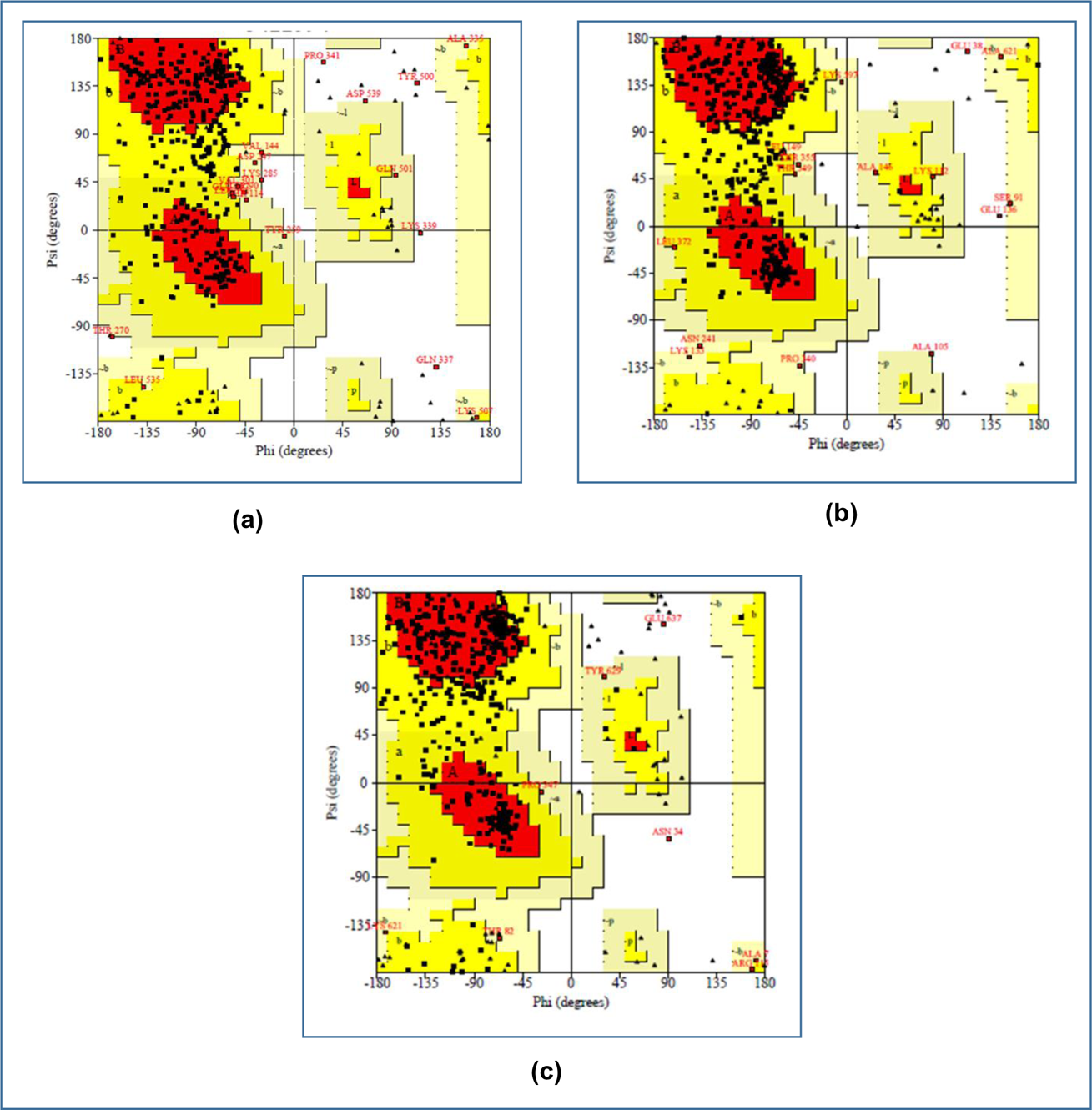
The Ramachandran plot analysis of the three vaccine constructs. (a) HV-1, (b) HV-2, (c) HV-3. The 3D structures of the constructed vaccines were refined using online refinement tool 3Drefine (http://sysbio.rnet.missouri.edu/3Drefine/) and validated by analyzing the Ramachandran plot, generated using the online tool PROCHECK (https://servicesn.mbi.ucla.edu/PROCHECK/).

### 3.11. Protein Disulfide Engineering

In protein disulfide engineering, disulfide bonds were generated for the 3D structures of the vaccine constructs. The DbD2 server identifies the pairs of amino acids that have the capability to form disulfide bonds based on the given selection criteria. In this experiment, we selected only those amino acid pairs that had bond energy value was less than 2.00 kcal/mol. The HV-1 generated 12 amino acid pairs that had the capability to form disulfide bonds. However, only 4 pairs were selected since they had bond energy, less than 2.00 kcal/mol: 19 Arg and 148 Gly, 21 Gly and 47 Arg, 177 Val and 162 Gly, 233 Pro and 251 Ser. HV-2 generated 21 pairs of amino acids that had the capability to form disulfide bonds, however, only 6 pairs were selected: 60 Glu and 67 Glu, 68 Phe and 118 Ala, 174 Ser and 217 Ser, 275 Gly and 319 Lys, 314 Ser and 325 Pro, 360 Gly and 369 Met. HV-3 generated 15 pairs of amino acids capable of forming disulfide bonds and only 3 pairs of the amino acids were selected: 191 Val and 248 Gly, 253 Ser and 290 Ser, 255 Asn and 282 Trp. The selected amino acid pairs formed the mutant version of the original vaccines in the DbD2 server (**Figure 08**).

**Figure 08.**
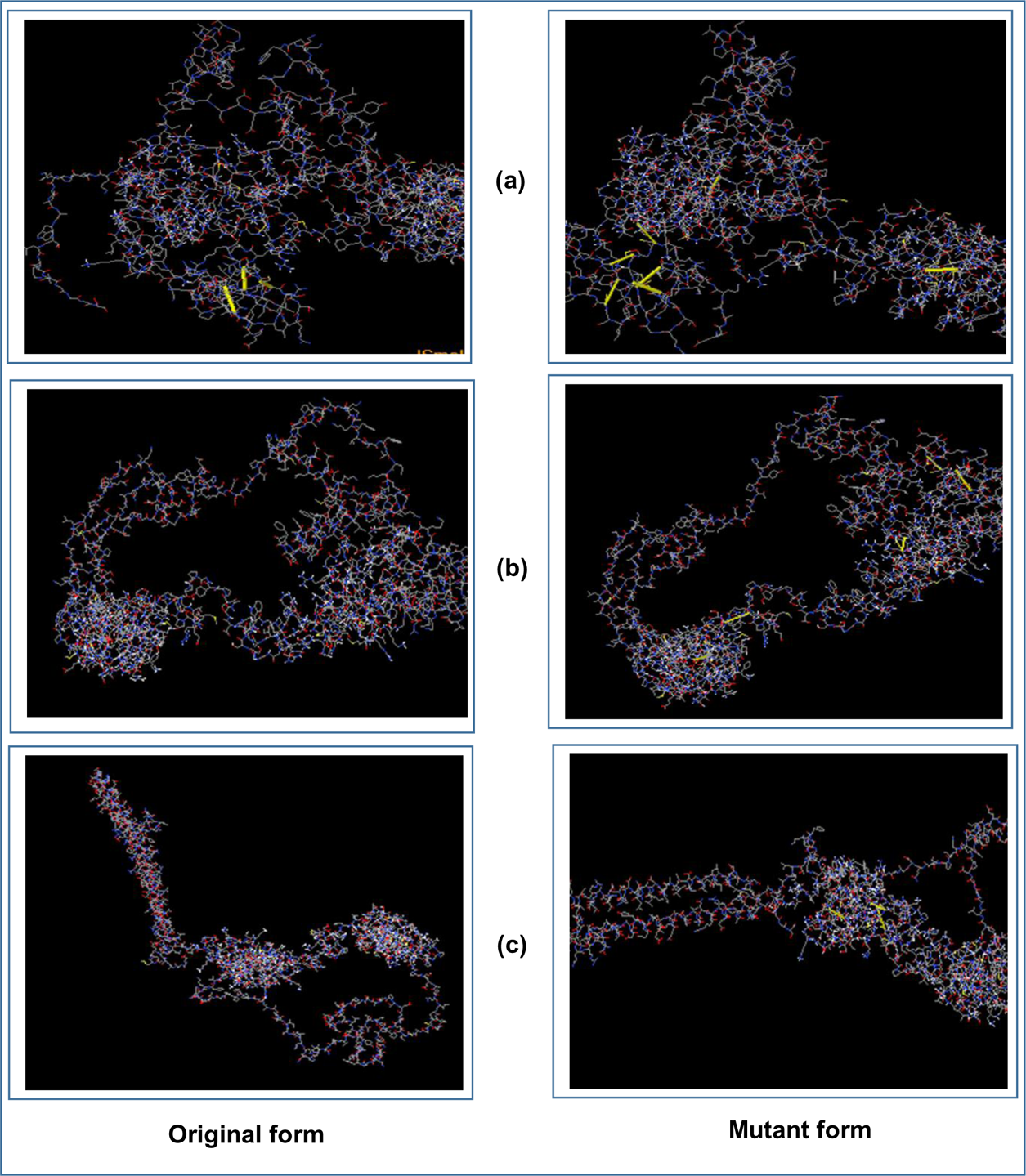
The disulfide engineering of the three vaccine constructs, both the original (left) and mutant (right) forms are shown. Here, (a) HV-1, (b) HV-2, (c) HV-3. The disulfide engineering was conducted using the online tool DbD2 server (http://cptweb.cpt.wayne.edu/DbD2/).

**Figure 09.**
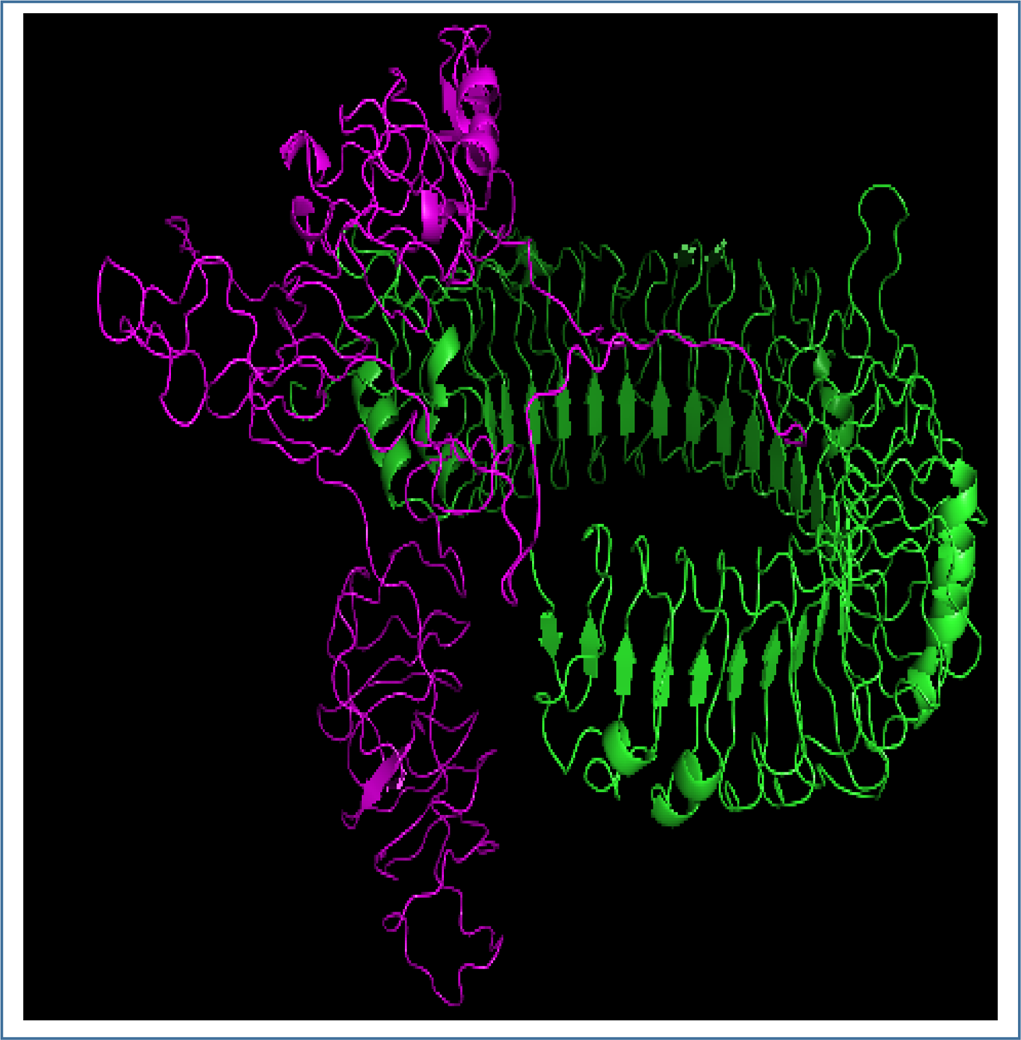
Figure showing the interaction between the TLR-3 and the selected vaccine HV-1. The interaction was visualized by PyMol tool. Here, the ligands is indicated by pink color and the receptor is indicated by green color.

### 3.12. Protein-Protein Docking Study

The protein-protein docking study was carried out to find out the best constructed Herpes vaccine. From analyzing the protein-protein docking, it was declared that HV-1 was the best constructed vaccine with the best and lowest scores in the MM-GBSA study and HawkDock study. However, when analyzed by ClusPro 2.0 and later refined and re-scored by the PRODIGY tool of HADDOCK server, HV-1 showed the best binding affinity, ΔG scores with DRB3*0202 (−17.1 kcal/mol), DRB3*0101 (−19.2 kcal/mol), DRB1*0401 (−21.2 kcal/mol) and TLR-3 (−21.9 kcal/mol). And when analyzed by PatchDock and FireDock servers, HV-1 showed best global energy scores with three MHC alleles: DRB5*0101 (−18.12), DRB1*0401 (−32.33), DRB1*0301 (−13.32) and TLR-3 (−10.66). Since HV-1 showed superior results in the protein-protein docking study, it was considered as the best vaccine construct among the three constructed vaccines (**Table 15**). The molecular dynamics simulation study and in silico codon adaptation studies were conducted on only the HV-1 vaccine.

**Table 15.**
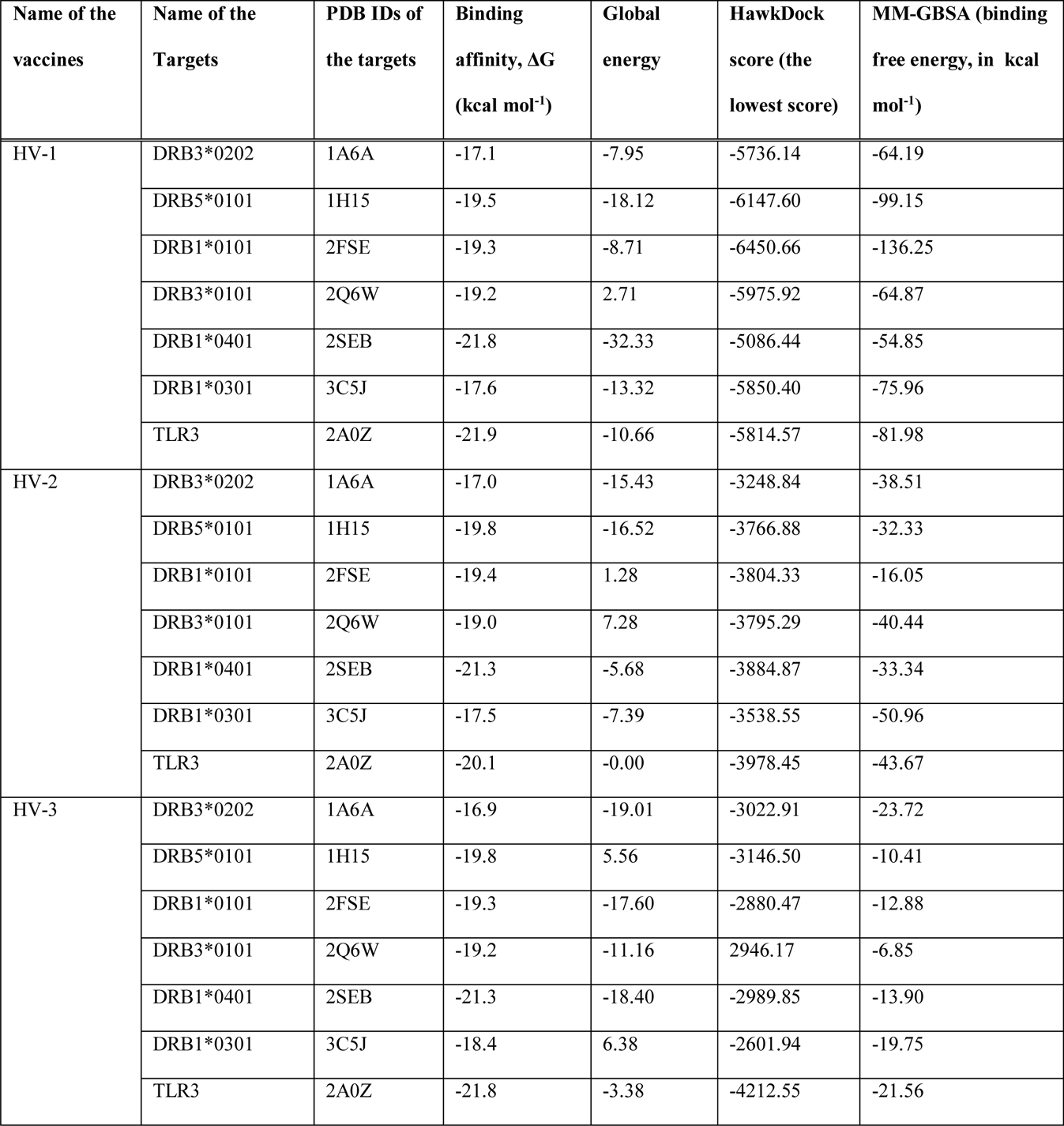
Results of the docking study of all the vaccine constructs.

### 3.13. Molecular Dynamics Simulation

**Figure 10** illustrates the results of molecular dynamics simulation and normal mode analysis (NMA) of HV-1-TLR-3 docked complex. The deformability graph of the complex indicates the peaks in the graphs which corresponds to the regions of the protein with possible deformability (**Figure 10b**). The B-factor graph of the complex gives easy visualization of the difference as well as comparison between the NMA and the PDB field of the docked complex (**Figure 10c)**. The eigenvalue of the complex is illustrated in **Figure 10d**. HV-1 and TLR-3 docked complex generated eigenvalue of 1.042621e-04. The variance graph indicates the individual variance by red colored bars and cumulative variance by green colored bars (**Figure 10e**). **Figure 10f** illustrates the co-variance map of the complex where the correlated motion between a pair of residues is indicated by red color, uncorrelated motion is indicated by white color and anti-correlated motion is indicated by blue color. The elastic map of the complex represents the connection between the atoms and darker gray regions correspond to the stiffer regions (**Figure 10g**) [67, 68, 69].

**Figure 10.**
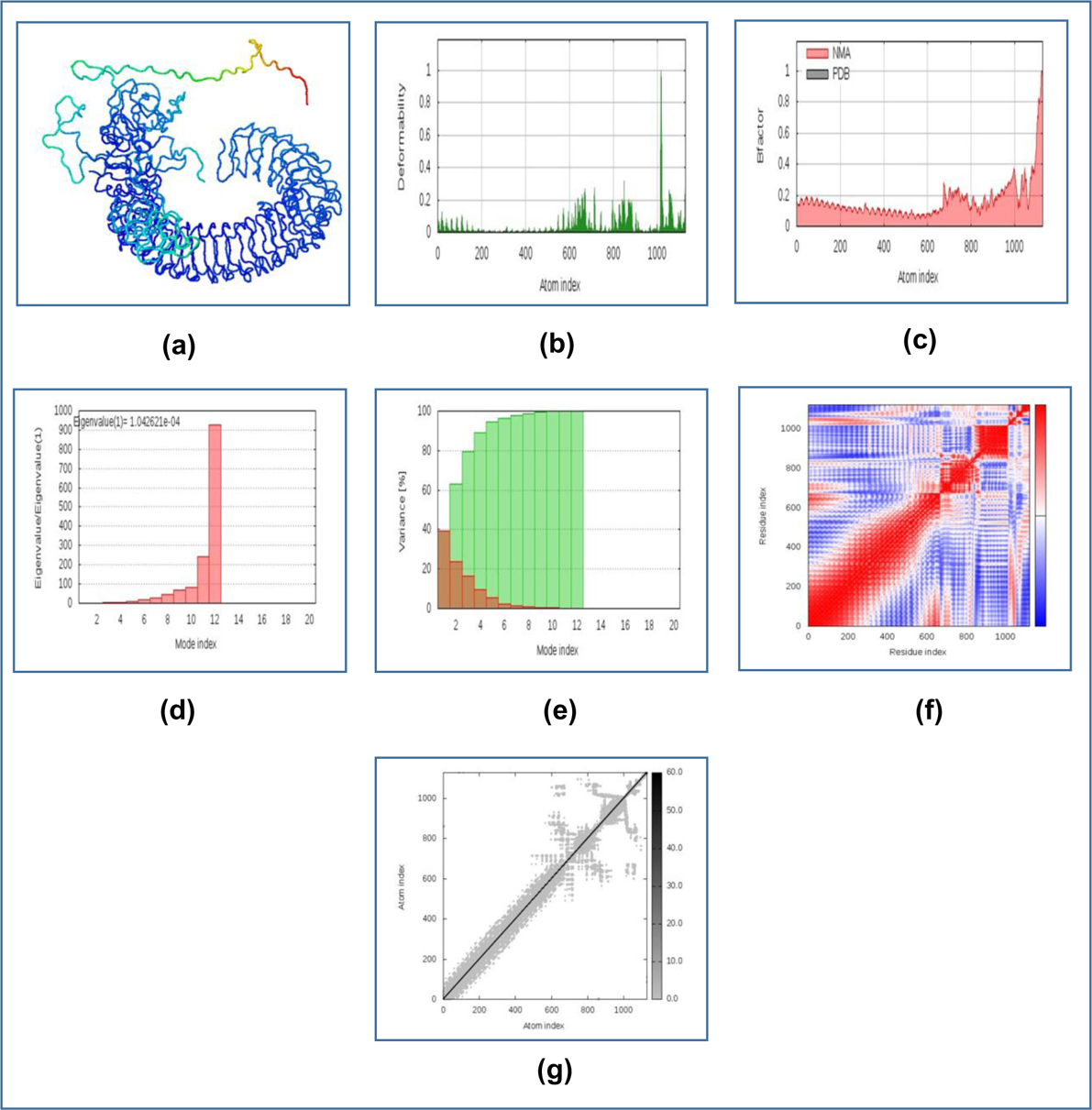
Figure displaying the results of molecular dynamics simulation study of HV-1 and TLR-3 docked complex. Here, (a) NMA mobility, (b) deformability, (c) B-factor, (d) eigenvalues, (e) variance (red color indicates individual variances and green color indicates cumulative variances), (f) co-variance map (correlated (red), uncorrelated (white) or anti-correlated (blue) motions) and (g) elastic network (darker gray regions indicate more stiffer regions).

### 3.14. Codon Adaptation and In Silico Cloning

For in silico cloning and plasmid construction, the protein sequences of the best selected vaccines were adapted by the JCat server (http://www.jcat.de/).

Since the HV-1 protein had 600 amino acids, after reverse translation, the number of nucleotides of the probable DNA sequence of HV-1 would be 1800. The codon adaptation index (CAI) value of 0.973 of HV-1 indicated that the DNA sequences contained higher proportion of the codons that are most likely to be used in the cellular machinery of the target organism *E. coli* strain K12 (codon biasness). For this reason, the production of the HV-1 vaccine would be carried out efficiently [74, 75]. The GC content of the improved sequence was 56.33%. The predicted DNA sequence of HV-1 was inserted into the pET-19b vector plasmid between the SgrAI and SphI restriction sites. Since the DNA sequence did not have restriction sites for SgrAI and SphI restriction enzymes, SgrA1 and SphI restriction sites were conjugated at the N-terminal and C-terminal sites, respectively, before inserting the sequence into the plasmid pET-19b vector. The newly constructed cloned plasmid would be 7372 base pair long, including the constructed DNA sequence of the HV-1 vaccine (the HV-1 vaccine DNA sequence also included the SgrAI and SphI restriction sites) (**Figure 11** and **Figure 12**).

**Figure 11.**
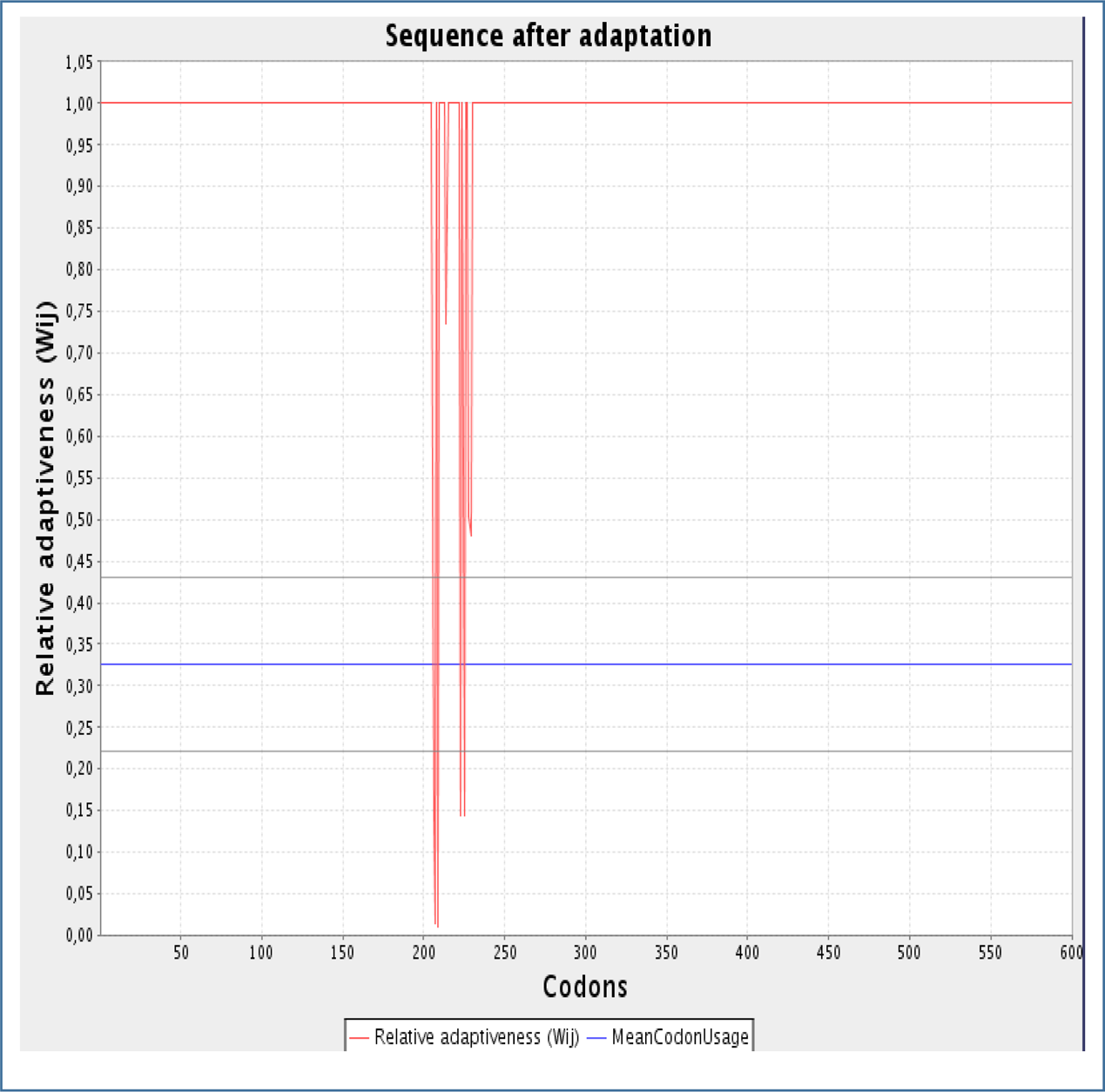
Figure showing the codon adaptation graphs of the HV-1 vaccine. The codon adaptation of the vaccine constructs were carried out using the server JCat (https://jcatbeauty.com/).

**Figure 12.**
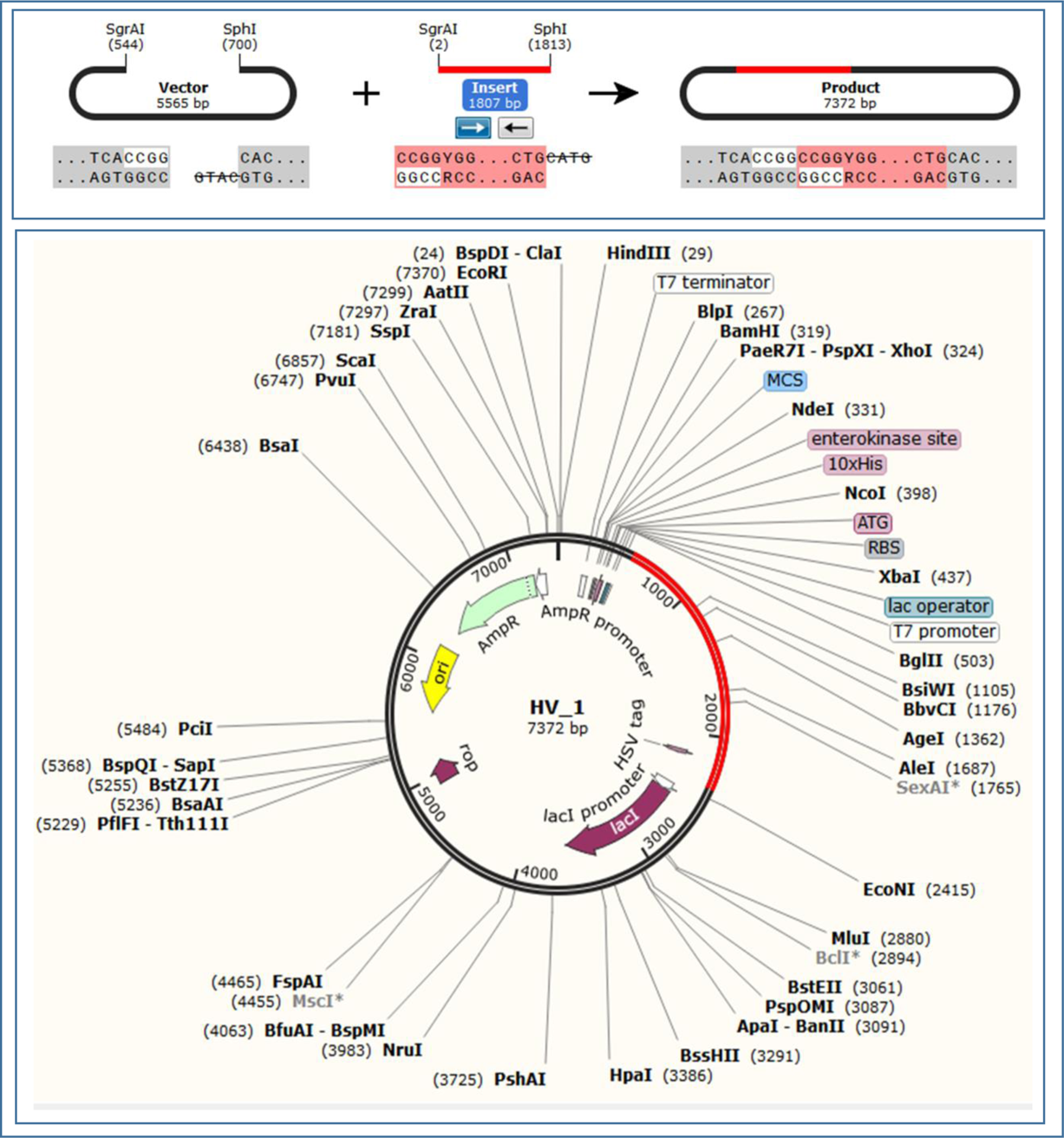
In silico restriction cloning of the HV-1 vaccine sequence in the pET-19b plasmid between the SgrAI and SphI restriction enzyme sites. The red colored marked sites contain the DNA inserts of the vaccines. The cloning was carried out using the SnapGene tool. The two newly constructed plasmids can be inserted into *E. coli* strain K12 for efficient vaccine production.

## 4. Discussion

Vaccine is one of the most important and widely produced pharmaceutical products. Millions of people and infants are getting vaccinated every year. However, the research and development processes of vaccines are costly and sometimes, it takes many years to develop a proper vaccine candidate against a particular pathogen. In recent times, various methods and tools of bioinformatics, immunoinformatics and reverse vaccinology are exploited for vaccine development, which save time and cost of the vaccine development process [76].

The current study was conducted to design and construct potential Herpes Simplex Virus (HSV), strain-17 vaccine. The target virus and its strain were identified by reviewing the NCBI database (https://www.ncbi.nlm.nih.gov/). Nine envelope glycoproteins were selected as targets for vaccine construction: envelope glycoprotein M, envelope glycoprotein H, envelope glycoprotein I, envelope glycoprotein E, envelope glycoprotein B, envelope glycoprotein L, envelope glycoprotein D, envelope glycoprotein K and envelope glycoprotein C. Envelope glycoprotein M plays important role in viral assembly [77]. The envelope glycoprotein H forms heterodimer with envelope glycoprotein L and both the envelope glycoproteins function in virus entry into the host cell. Moreover, envelope glycoprotein B, together with envelope glycoprotein H and envelope glycoprotein K, aids in the fusion events of the virion envelope with the outer nuclear membrane [78, 79, 80]. The main functions of envelope glycoprotein E and envelope glycoprotein I are to spread the virus from cell to cell as well as the fusion of the cells [81, 82, 83]. The envelope glycoprotein D also helps the virus to enter into the host cell [84]. The envelope glycoprotein C aids in the adsorption of the virion to the host cell surface [85]. Since, the glycoproteins mainly function in the infection process of the HSV, for this reason, targeting these glycoproteins could be potential strategy for vaccine construction.

After selecting the proteins, the antigenicity of the proteins were determined by VaxiJen V2.0 server. The proteins that were found to be antigenic, were selected for further analysis. Three glycoproteins, envelope glycoprotein E, envelope glycoprotein B and envelope glycoprotein D were found to be antigenic and for this reason, they were selected for further analysis. The various physicochemical properties like number of amino acids, molecular weight, theoretical pI, extinction co-efficient, instability index, aliphatic index, GRAVY were determined by ProtParam (https://web.expasy.org/protparam/) server. The three proteins performed quite similarly in the physicochemical property test.

Two of the main cells that function in immunity are the T lymphocytic cell and B lymphocytic cell. After recognized by an antigen presenting cell or APC (like macrophage, dendritic cell etc.), the antigen is presented through the MHC class-II molecule present on the surface of the APC, to the helper T cell. Since, the helper T cell contains CD4+ molecule on its surface, it is also known as CD4+ T cell. After activated by the antigen presenting cell, the T-helper cell then activates the B cell and cause the production of memory B cell and antibody producing plasma B cell. The plasma B cell produce a large number of antibodies and the memory B cell functions as the immunological memory. However, macrophage and CD8+ cytotoxic T cell are also activated by the T-helper cell, that destroy the target antigen [86, 87, 88, 89, 90]. The possible T cell and B cell epitopes of the selected viral proteins were determined by the IEDB (https://www.iedb.org/) server. The IEDB server generates and ranks the T cell epitopes based on their antigenicity scores (AS) and percentile scores. The top ten MHC class-I epitopes were taken for analysis. However, since more than one epitope showed similar AS and percentile score, two epitopes were selected from each of the AS and percentile score categories. However, based on analyzing the lengths of the B-cell epitopes, five epitopes were selected for further analysis. The transmembrane topology of the selected eiptopes were predicted to determine whether they would be present at the exterior or interior of the viral envelope.

Antigenicity is defined as the ability of a foreign substance to act as antigen and activate the T cell and B cell responses, through their antigenic determinant portion or epitope [91]. The allergenicity of a substance is defined as the ability of that substance to act as allergen and induce potential allergic reactions within the body [92]. On the other hand, when designing a vaccine, epitopes of that remain conserved across various strains, are given much priority than genomic regions that are highly variable among the strains since the conserved epitopes of protein(s) provide broader protection across various strains and sometimes even species [93]. When selecting the best epitopes for vaccine construction, some criteria were maintained: the epitopes should be highly antigenic, so that they can induce high antigenic response, the epitopes should be non-allergenic in nature, for this reason, they would not be able to induce any allergenic reaction in an individual and the epitopes should be non-toxic. The epitopes with 100% conservancy and over 50% minimum identity were selected for vaccine construction, so that, the conserved epitopes would be able to provide protection against various strains. The conservancy analysis of the epitopes were carried out using Human Herpes Simplex Virus-1 strain-F and Human Herpes Simplex Virus-2 strain-HG52, for comparison, so that the conservancy analysis across the strains as well as species could be carried out efficiently. Total 18 T-cell epitopes (six epitopes from each of the proteins) and 9 B-cell epitopes were selected for vaccine construction. Moreover, the cluster analysis of the MHC class-I alleles and MHC class-II alleles were also carried out to determine their relationship with each other and cluster them functionally based on their predicted binding specificity [94].

In the next step, the protein-peptide docking was carried out between the selected epitopes and the MHC alleles. The MHC class-I epitopes were docked with the MHC class-I molecule (PDB ID: 5WJL) and the MHC class-II epitopes were docked with the MHC class-II molecule (PDB ID: 5JLZ). The protein-peptide docking was performed to determine the ability of the epitopes to bind with their respective MHC molecule. After 3D structure generation of the epitopes, the docking was carried out by PatchDock and FireDock servers. YTLSVGDIK, ASANASVER, KIAGWHGPK, PSWVSLMPPKQV, TMYYKDVTVSQV and QAYQQGVTVDSI generated the best scores in the protein-peptide docking. However, all of the selected T-cell epitopes showed significant interaction capability with their respective targets.

After successful protein-peptide docking, the vaccine construction procedure was carried out. Three different vaccines were constructed, that differ from each other based on their adjuvant sequences. The three vaccine constructs were designated as: HV-1 (600 amino acids long), HV-2 (685 amino acids long) and HV-3 (714 amino acids long).

After the vaccine construction was performed, the antigenicity, allergenicity and physicochemical property analysis were carried out. All the vaccine constructs were proved to be antigenic, as well as non-allergenic, for this reason, they should not cause any allergenic reaction within the body, however, all of them should be able to induce high immunogenic response. The extinction coefficient corresponds to the amount of light, that is absorbed by a compound at a certain wavelength [95, 96]. HV-3 had the highest extinction co-efficient of 74566.83 M-1 cm-1. The aliphatic index of a protein referes to the relative volume occupied by the aliphatic amino acids in the side chains like alanine, valine etc. [97]. HV-2 had the highest aliphatic index among the vaccine constructs, although HV-3 had aliphatic index that is very near to the aliphatic index of HV-2. All the three vaccine constructs had the similar half-life of 1 hour in the mammalian cell culture and all of them were stable. All of the constructs had quite similar theoretical pI. Quite similar performances were observed by the three vaccine constructs in the physicochemical property analysis.

Two online servers, PRISPRED (http://bioinf.cs.ucl.ac.uk/psipred/) and NetTurnP v1.0 (http://www.cbs.dtu.dk/services/NetTurnP/), were used for protein secondary structure determination. HV-3 had the highest percentage of amino acids in the alpha formation and lowest percentage of amino acids in the beta-sheet formation as well as in coil structure formation. The 3D structures were generated by the server RaptorX (http://raptorx.uchicago.edu/) server and with the lowest p-value of 8.91e-05, HV-2 showed the best performance on the protein 3D structure generation. However, all of the proteins had 3 domains. After 3D structure generation, the 3D structures were refined by online tool 3Drefine (http://sysbio.rnet.missouri.edu/3Drefine/) and validated by PROCHECK (https://servicesn.mbi.ucla.edu/PROCHECK/) server. HV-3 had the best performance in the 3D structure refinement and validation study with 74.7% of the amino acids in the most favored regions and 23.7% of the amino acids in the additional allowed regions. After validation of the 3D protein structures, the disulfide engineering of the vaccine constructs were performed using Disulfide by Design 2 v12.2 (http://cptweb.cpt.wayne.edu/DbD2/) server and the amino acid pairs with binding energy value less than 2.0 kcal/mol, were chosen for disulfide bond formation. With six pairs of amino acids, that had binding energy less than 2.0 kcal/mol, it can be considered that, HV-2 showed the best performance in the protein disulfide engineering study.

In the next step, the protein-protein docking was carried out using the vaccine constructs as ligands and various MHC alleles as receptors. The docking experiment was carried out to determine the best vaccine construct among the three vaccines. All the vaccines successfully bound with their target receptors. However, from the docking experiment, it was revealed that, HV-1 was the best vaccine among the three vaccine constructs. For this reason, HV-1 vaccine construct was used for molecular dynamics simulation and in silico cloning. The molecular dynamics simulation study showed that, the HV-1-TLR-3 complex had very low chance of deformability. This was further confirmed by the deformability graph of 10b where the spikes in the graphs rarely reaches to the maximum value. The eigenvalue of the complex of 1.042621e-04 also pointed to the fact that the HV-1-TLR-3 docked complex should be quite stable and should have relatively less chance of deformability. The **Figure 10f** illustrates the correlated motion of the amino acid residue pairs in red color and according to the figure, the complex showed a large amount of amino acid pairs in the correlated motion. Finally the HV-1 vaccine was reverse transcribed into the possible DNA sequence for cloning into the *E. coli* strain K12. Before in silico cloning procedure was performed, codon adaptation of the HV-1 vaccine was carried out, where the CAI value of 0.973 was achieved, which reflects that the newly adapted sequence had high degree of codons that are mostly likely to be used by *E. coli* strain K12. Moreover, good GC content of 56.33% was also achieved. Finally, the adapted DNA sequence was inserted into the pET-19b plasmid and the plasmid with the HV-1 insert was now 7372 base pairs long. This newly generated plasmid with the target DNA sequence can be inserted into the *E. coli* strain K12 for efficient production of the HV-1 vaccine.

However, more in vitro and in vivo researches should done to finally confirm the safety, efficacy and potentiality of the predicted vaccines.

## 5. Conclusion

Herpes Simplex Virus is one of the most infectious and sexually transmitted diseases in the world. However, only the HSV-2 is transmitted sexually. The other type of Herpes virus, HSV-1, transmits though oral-oral contact. With very high infection rate, HSV can be considered as one of the fatal viruses in the world. However, no vaccines are yet discovered with satisfactory results, to control the infection of HSV. In this study, a possible subunit vaccine to fight against HSV-1, strain-17, was designed using various tools of bioinformatics, immunoinformatics and vaccinomics. In this study, first the potential proteins of the viral structure were first identified. Next, the potential epitopes were identified through robust processes and these epitopes were used for vaccine construction. Since only the highly antigenic and at the same time, non-allergenic epitopes were selected, the constructed vaccines should confer very strong antigenic response, however, with no allergenicity, the constructed vaccines should be safe to be administered. Three possible vaccines were constructed and by conducting the docking experiments, one best vaccine construct was identified. Later, the molecular dynamics simulation study and codon adaptation as well as in silico cloning were performed for large scale production of the vaccines. The vaccine should provide better immunity towards the HSV-1, strain-17 as well as the HSV-1 strain-F and the HSV-2, strain-HG52, due to its conservancy across the strains and species. However, wet lab researches should be carried out on the findings of this experiment to finally confirm the safety, efficacy and potentiality of the vaccine constructs. Hopefully, this research will raise interests among the scientists of the respective field.

## 6. Acknowledgements

Authors are thankful to Swift Integrity Computational Lab, Dhaka, Bangladesh, a virtual platform of young researchers, for providing the tools.

## 7. Conflict of interest

Bishajit Sarkar declares that he has no conflict of interest. Md. Asad Ullah declares that he has no conflict of interest.

